# Single cell discovery of m^6^A RNA modifications in the hippocampus

**DOI:** 10.1101/2023.12.06.570314

**Authors:** Shuangshuang Feng, Maitena Tellaetxe-Abete, Yujie Zhang, Yan Peng, Han Zhou, Erika Larrea, Liang Xue, Li Zhang, Magdalena J. Koziol

**Affiliations:** State Key Laboratory of Cognitive Neuroscience and Learning, Beijing Normal University, Beijing, 100875, China; Chinese Institute for Brain Research, Beijing, 102206, China; Research Unit of Medical Neurobiology, Chinese Academy of Medical Sciences, Beijing, 102206, China; Intelligent Systems Group, Computer Science Faculty, University of the Basque Country, Donostia/San Sebastian, 20018, Spain; Peking University, Beijing, 100871, China; Tsinghua University, Beijing, 100084, China

## Abstract

*N*^6^-methyladenosine (m^6^A) is a prevalent and highly regulated RNA modification essential for RNA metabolism and normal brain function. It is particularly important in the hippocampus, where m^6^A is implicated in neurogenesis and learning. Although extensively studied, its presence in specific cell types remain poorly understood. We investigated m^6^A in the hippocampus at the single cell level, revealing a comprehensive landscape of m^6^A modifications within individual cells. Our data also identifies transcripts which have high m^6^A density and are associated with brain diseases. Our data suggests that m^6^A containing transcripts might be of particular importance for *Camk2a* neurons. Overall, this work provides new insights into the molecular mechanisms underlying hippocampal physiology and lays the foundation for future studies investigating the dynamic nature of m^6^A RNA methylation in the healthy and diseased brain.

## INTRODUCTION

The hippocampus plays a critical role in learning and memory formation (Bird and Burgess 2008). Understanding the molecular mechanisms underlying hippocampal function is therefore of great importance in the field of neuroscience. N^6^-methyladenosine (m^6^A) RNA methylation, a prevalent and highly regulated epitranscriptomic modification, has emerged as a crucial regulatory mechanism in various biological processes, including neurodevelopment and learning (Li et al. 2017; Livneh et al. 2020; Jiang et al. 2021; Yu et al. 2021). Since abnormal m^6^A levels are associated with human neurological diseases (Lv et al. 2023), studying m^6^A in the brain is of particular interest.

The dynamic m^6^A modification is catalyzed by a multi-protein complex involving the methyltransferase METTL3 (methyltransferase-like 3), which installs a methyl group onto adenosine residues of RNA transcripts (Liu et al. 2014). m^6^A modifications can be removed by other enzymes, such as the dioxygenase FTO (Fat mass and obesity associated) and ALKBH5 (Li et al. 2017). The recognition and interpretation of m^6^A-modified transcripts are mediated by reader proteins, among which YTHDF2 (YTH N^6^-methyladenosine RNA binding protein 2) has been extensively studied(Wang et al. 2014; Du et al. 2016).

While the significance of m^6^A RNA methylation in various tissues has been extensively investigated, its precise roles and impact in specific cell types of the brain, such as the hippocampus, are poorly understood. Brain regions such as the hippocampus consist of many different specialized cells. Previous studies have demonstrated the abundant presence of m^6^A modifications in hippocampal cells (Zhou et al. 2023). However, it is currently unknown which cell types and transcripts have m^6^A, and with which density.

Studies utilizing *Mettl3*, *Ythdf2* and *Fto* knockout mice have shown impaired hippocampal-dependent learning and memory (Li et al. 2017; Engel et al. 2018; Livneh et al. 2020; Zhuang et al. 2023). In contrast, overexpression of *Mettl3* significantly enhances long-term memory consolidation(Zhang et al. 2018b; Jiang et al. 2021). Chang *et al*. demonstrated distinct levels of m^6^A modification in different cellular subtypes of the cerebellum and cortex, suggesting potential cell-type-specific functions(Chang et al. 2017).

To comprehensively understand the landscape of m^6^A RNA methylation in the hippocampus and its relevance to specific cell types, it is crucial to investigate this modification at the single cell level. Single cell sequencing technologies have revolutionized our ability to uncover cellular heterogeneity and identify distinct subpopulations within complex tissues (Regev et al. 2017). Applying these technologies to the study of m^6^A modifications in the hippocampus will provide unprecedented insights into cell-type-specific roles of m^6^A in hippocampal function facilitating the development of future targeted therapeutics.

In this study, we aim to address the knowledge gap by performing single cell RNA sequencing to profile m^6^A RNA methylation patterns in individual cells of the hippocampus. A myriad of different m^6^A sequencing detection methods exist, such as m^6^A RNA immunoprecipitation (RIP)(Dominissini et al. 2012; Meyer et al. 2012) coupled with next generation sequencing technologies, or m^6^A-SEAL(Wang et al. 2020) or MAZTER-seq (Garcia-Campos et al. 2019b). However, all of them require high amounts of RNA input, currently making them incompatible with single cell approaches in regular somatic cells(Ke et al. 2015; Linder et al. 2015; Garcia-Campos et al. 2019a; Zhang Zhang 2019; Shu et al. 2020; Wang et al. 2020; Li et al. 2023; Yao et al. 2023). Due to the limited amount of RNA input in smaller somatic cells, such as found in the brain, the use of m^6^A-RIP to detect m^6^A at single cell level seems currently not feasible. Deamination adjacent to RNA modification targets sequencing (DART-seq) is another method that facilitates m^6^A detection(Meyer 2019). It utilizes the expression of a fusion protein APOBEC1-YTH, in which the APOBEC1 protein is fused to the YTH m^6^A-binding domain of YTHDF2. In this construct, the YTH domain lures APOBEC1 to the close vicinity of m^6^A sites, where APOBEC1 deaminates cytidine into uracil (Fig. 1A)(Meyer 2019). By identifying C-to-U editing events that correspond to C-to-T mutations in sequencing data, adjacent m^6^A sites can be identified. 97% of these sites disappeared in methylase METTL3-depleted cells, illustrating the specificity of this method(Meyer 2019). The feasibility of this single cell DART-seq (scDART- seq) approach has recently also been demonstrated using a relatively homogenous HEK293T cell line (Tegowski et al. 2022). Here, we use this approach to identify how m^6^A is distributed in different cell types of the hippocampus to gain a comprehensive understanding of the cell-type-specific distribution and characteristics of m^6^A modifications.

**Figure 1.**
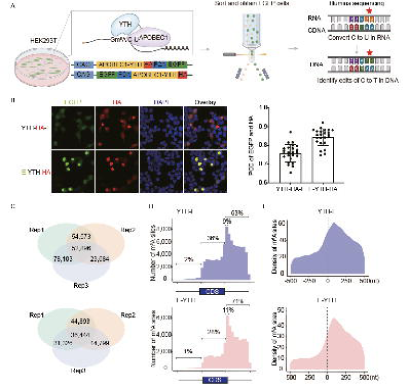
Improved m^6^A DART-seq detection in HEK293T cells. (A) Schematic diagram of m^6^A detection with DART-seq in cultured cells. The YTH protein domain binds to m^6^A. When bound to APOBEC1, the APOBEC1 protein converts C-to-U in the vicinity of m^6^A. This results in a C-to-T mutation in cDNA. C-to-T mutations detected by RNA sequencing are indicative of m^6^A RNA modifications. (B) Left: Immunofluorescence (IF) of HEK293T cells transfected with *Apobec1-Yth-HA-Egfp* (*Yth-HA-E*) or *Egfp-Apobec1-Yth-HA (E-YTH-HA*). Scale bar, 20μm, representative images are shown. Right: Quantification of EGFP and HA overlap. PCC: Pearson’s Correlation Coefficient. (C) Number of C-to-T editing events identified in each bulk DART-seq HEK293T cell replicate for *Yth-E* and *E-Yth* plasmids. Editing events identified in at least 2 replicates were considered for downstream analyses. The data was obtained following *Apobec1-Yth-Egfp* or *Egfp-Apobec1-Yth* transfection and EGFP FACS sorting. n=3, Rep: separately cultured replicate. (D) Metagene analysis showing m^6^A site counts along transcripts for *Yth-E* and *E-Yth* DART-seq results. 9% of all m^6^A sites occur in the first 10% of the 3’ UTR following the TTS for YTH-E, and 11% for E-YTH, respectively. Shown percentage indicates number of m^6^A sites upstream, within and downstream of coding sequence (CDS). (E) Metagene analysis showing m^6^A density 500nt 5’ and 500nt 3’ from stop codon (0nt) for YTH-E and E-YTH.

## RESULTS

### Confirmation of m^6^A detection with DART-seq

To be able to apply the DART-seq approach in cells of the mouse hippocampus *in vivo*, we subcloned *Apobec1-Yth* with an HA tag into an adeno-associated virus (AAV) plasmid under control of a chicken β actin (CAG) promoter. As controls, we also generated *Apobec1*-*Yth^mut^*that can no longer bind m^6^A, as well as *Apobec1* only. Through AAV transduction, these proteins can then be expressed in different cell types of the mouse brain(Negrini et al. 2020b). To be able to isolate and analyze only successfully AAV transfected cells, a P2A sequence followed by *Egfp* was added to the C-terminal. This should enable selection while avoiding potential interference with APOBEC1-YTH function. We argued that FACS selection for DART- seq positive cells is essential, as cells and transcripts that have not been exposed to APOBEC1- YTH sufficiently could result in many false negatives and therefore in higher m^6^A heterogeneity. To test our *Apobec1*-*Yth*-*HA*-*P2A*-*Egfp* (*Yth-E*) plasmid, we transfected *Yth*-*E* into human HEK293T cells (subsequently referred to as YTH-E sample). Although *Egfp* and *Apobec1*-*Yth*- HA were co-expressed, the reporter signal of enhanced green fluorescent protein (EGFP) was weak (Fig. 1B). Similarly, when *Yth*-*E* was transfected into the hippocampus, an EGFP signal could not be easily detected with FACS sorting (Fig. 1B; Supplemental Fig. S1A). We argued that this low signal is because *Egfp* is located at the C-terminal of one long RNA transcript(Zhu et al. 2023). We therefore changed the order of the expression cassette on either side of P2A, moving the *Egfp* to the N-terminus, to make the *Egfp*-*P2A*-*Apobec1*-*Yth*-*HA* (*E-Yth*) and *Egfp*- *P2A*-*Apobec1*-*Yth^mut^*-*HA* (*E*-*Yth^mut^*) plasmids. Indeed, *E*-*Yth* transfected HEK293T cells (subsequently referred to as E-YTH sample) resulted in an increased EGFP signal and improved the correlation with *Apobec1*-*Yth*-*HA* (Fig. 1B). We also detected EGFP in both cytoplasm and nuclei for E-YTH and E-YTH^mut^ samples (Supplemental Fig. S1B), suggesting that both could be used for single cell sequencing.

To evaluate our E-YTH construct further, we transfected *E-Yth* and *Yth-E* into HEK293T cells, followed by FACS sorting and RNA sequencing (RNA-seq) (Fig. 1A). As controls, we processed *E-Yth^mut^*, *Yth^mut^-E*, *E-Apobec1*, *Apobec1*-*E* and mock transfected cells. We only considered C-to-U editing sites that were identified in at least 2 replicates, such as 209,256 C-to-U editing sites in YTH-E and 127,462 in E-YTH samples (Fig. 1C; Supplemental Table S1) and did similar analysis for control samples (Supplemental Fig. S1C; Supplemental Table S1). To identify m^6^A sites in E- YTH and YTH-E, we eliminated background C-to-U editing events detected in APOBEC1 only, YTH^mut^ and mock transfected cells (more details in Methods). We identified 106,827 m^6^A sites that occur in transcripts in YTH-E and 54,395 in E-YTH (Supplemental Table S2). In both cases we found m^6^A to be enriched in the 3’UTR region, 63% in YTH-E and 71% in E-YTH, confirming previous m^6^A findings (Fig. 1D,E; Supplemental Table S3)(Dominissini et al. 2012; Meyer et al. 2012). 9% of all m^6^A sites in YTH-E and 11% in E-YTH occur in the first 10% of the 3’ UTR (Fig. 1D,E; Supplemental Table S3). Overall, YTH-E and in E-YTH confirm previous m^6^A findings. Although more m^6^A sites have been identified in YTH-E than in E-YTH, the distribution of m^6^A sites is more defined in YTH-E (Fig. 1D; Supplemental Fig. S1D). Not only has E-YTH a stronger EGFP signal, facilitating E-YTH isolation in the mouse hippocampus, but it shows higher correlation between EGFP signal and APOBEC1-YTH presence, providing more defined m^6^A distribution patters with less background (Fig. 1B,D). Overall, we decided to use E-YTH with the corresponding controls for our hippocampal studies.

### Detection of m^6^A with bulk RNA-seq in mouse hippocampus

Liquid Chromatography coupled with tandem mass spectrometry (LC-MS/MS) of 3-month-old adult mouse brains showed that mRNAs in hippocampi have higher levels of m^6^A than in mRNAs of the cortex, thalamus, or cerebellum. In hippocampal mRNAs, we found that m^6^A occurs 1 per 1,000 unmodified adenosines (Supplemental Fig. S2A). To identify m^6^A within hippocampal mRNA sequences, we tested if E-YTH can facilitate m^6^A detection *in vivo*. Thus, we packaged *E-Yth* and controls *E-Yth^mut^* and *E-Apobec1* into AAV viruses for local hippocampal stereotaxic injection (Fig. 2A). After verifying that the AAV viruses were successfully expressed in the hippocampus (Fig. 2B), EGFP positive cells were collected by FACS sorting and processed for RNA-seq (Fig. 2A). Hippocampi wild type samples were also processed as additional controls. 2,672 and 2,272 C-to-U editing sites were identified in all 3 replicates of E-YTH and E- YTH^mut^, respectively. Only a few hundred C-to-U editing sites overlap between 2 replicates alone, while more are common between all 3 replicates, demonstrating high reproducibility between replicates (Fig. 2C; Supplemental Fig. S2B; Supplemental Table S4). To identify m^6^A sites, we only considered C-to-U editing events that occur in at least 2 out of 3 replicates (Fig. 2C; Supplemental Table S4) and removed background events from E-YTH, such as editing events detected in wild type, APOBEC1 and E-YTH^mut^ (see Methods; Supplemental Fig. S2B; Supplemental Table S4). In total, we identified 1,578 edits in E-YTH transcripts, but only 82 edits in E-YTH^mut^ controls (Supplemental Table S5). To further verify the feasibility of m^6^A detection with E-YTH in the mouse hippocampus, we detected 8,220 m^6^A regions that correspond to 5,431 genes with m^6^A-RIP (Supplemental Fig. S2C; Supplemental Table S6). 32% of E-YTH m^6^A sites were also detected and corroborated with m^6^A-RIP (Supplemental Fig. S2D; Supplemental Table S7). In accordance to previous studies using m^6^A-RIP and DART-seq in cells(Meyer et al. 2012), our E-YTH identified m^6^A are enriched in the 3’ region (80.8%) (Fig. 2D; Supplemental Fig. S2E), particularly near the TTS (Fig. 2E). We detected that C-to-U editing occurs 118bp downstream of the TTS site (Fig. 2E). To determine how often m^6^A occurs at a particular site on multiple transcript copies transcribed from one gene (here referred to as RNA replicates), we calculated the mutation per read ratio (m/k) for each editing site but excluded sites with fewer than 10 reads (see Methods). A m/k ratio of 1 indicates that all copies of a particular RNA have m^6^A, notably at the same site, while a m/k ratio of 0.25 stipulates that 25% of them have m^6^A in common. Our bulk sequencing data shows that m^6^A density at each position is low. Although there are a few transcripts that have m^6^A on all their RNA copies (m/k=1), most transcripts have m^6^A in less than 10% of their transcripts (Fig. 2G; Supplemental Fig. S1D). The bulk sequencing data suggests that m^6^A, although functionally important in the hippocampus (Zhang et al. 2018a; Du et al. 2021; Yin et al. 2023), is mostly heterogenous at specific sites. However, certain cells have sites where m^6^A occurs on many RNA replicates. To investigate this, we next applied our system to detect m^6^A sites on a single cell level in the mouse hippocampus.

**Figure 2.**
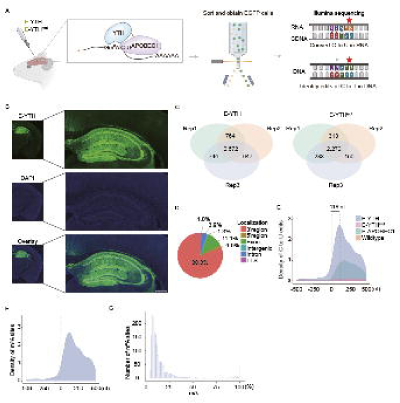
Detection of m^6^A with bulk DART-seq in mouse hippocampus. (A) Schematic diagram of DART-seq in the mouse brain. The *Egfp-Apobec1-Yth* is packaged into AAV viruses to infect brain cells. EGFP positive cells are isolated from the hippocampus and processed for C-to-U edit and m^6^A site identification. (B) Confocal image of mouse hippocampus after AAV infection. Representative image of E-YTH is shown. Half-brain image: Scale bar, 1mm. Hippocampus image: Scale bar, 400μm. (C) Number of overlapping C-to-U editing events identified by RNA-seq in hippocampus following *Egfp-Apobec1-Yth* and *Egfp-Apobec1-Yth^mut^* AAV virus injection and EGFP FACS sorting. Editing events identified in at least 2 replicates were considered for downstream analyses. n=3, Rep: biological replicates from different animals. (D) Pie chart showing m^6^A localization identified by *E-Yth* in mouse hippocampus. TTS: transcription termination site. (E) Metagene analysis showing C-to-U edit scaled density 500nt 5’ and 500nt 3’ from stop codon (0nt) in E-YTH, E-YTH^mut^, E-APOBEC1 and wild type samples. The peak value for E-YTH are 2,716 editing events and it occurs 118nt downstream of the stop codon. (F) Metagene analysis showing m^6^A density 500nt 5’ and 500nt 3’ from stop codon (0nt). m^6^A sites were obtain after eliminating background from E-YTH editing sites. (G) Histogram of m^6^A site counts over mutation per read (m/k) ratio. A minimum threshold of 5% was applied.

### Single cell sequencing of E-YTH transfected hippocampus

We injected AAV containing *E-Yth* or *E-Yth^mut^* into mouse hippocampi of 3-month-old mice (Fig. 3A). Since many cells in the hippocampus are fragile and vary greatly in shape and size, we used our optimized protocol for dissociating the mouse hippocampus into single cells (see Methods). We next FACS sorted for EGFP to only consider cells successfully transduced with *E- Yth* or *E-Yth^mut^*. We argued that such selection is essential, as only by eliminating cells that do not express E-YTH or E-YTH^mut^ sufficiently high enough, we can exclude m^6^A false negatives that could lead to overestimations of m^6^A heterogeneity in RNA replicates. After stringent EGFP selections, cells were then loaded into a microwell (Rhapsody, BD) for barcoding, followed by library generation and subsequent high-throughput sequencing (Fig. 3A; Supplemental Fig. S3A). We confirmed that our E-YTH and E-YTH^mut^ single cell libraries were of high quality and had sufficient reads (see Methods; Supplemental Fig. S3B; Supplemental Table S8). 11,561 cells with barcodes and UMIs were identified in E-YTH and 16,243 in E-YTH^mut^ samples (Supplemental Table S8). We then performed clustering on the integrated E-YTH and E-YTH^mut^ single cell datasets (Fig. 3B). The Uniform Manifold Approximation and Projection (UMAP) dimension reduction method was applied which identified 28 clusters. Except for cluster 23, which is entirely missing from our E-YTH data, all other clusters shows similar cell numbers between E-YTH and E-YTH^mut^ (Fig. 3C; Supplemental Fig. S3C). The missing of cluster 23 was unexpected, as the number of genes detected in all other clusters is very similar, with an average of 2,800 for E-YTH and 2,373 for E-YTH^mut^ (Fig. 3D). Also, few transcriptional changes were detected in E-YTH versus E-YTH^mut^ single cell data (Supplemental Fig. S4A-B). Since clusters 21, 22, 24-27 have fewer cell numbers than cluster 23 of E-YTH^mut^ (Fig. S3C), the absence of cluster 23 in E-YTH cannot be attributed to not enough sequencing depth. Thus, the absence of cluster 23 seems E-YTH specific. Automatic gene annotation identified cluster 23 as neurons (Supplemental Fig. S4C). Although there were other genes that are specifically enriched in cluster 23 and are not present in other clusters of the E-YTH^mut^ data (Supplemental Table S10), *Camk2a* draw our attention as it is a well characterized gene expressed in a subgroup of excitatory neurons of the hippocampus and cortex(Yasuda et al. 2022). Also, specific METTL3 depletion in CAMK2A expressing cells in the mouse hippocampus was previously shown to reduce long-term memory consolidation, highlighting the functional and biological importance of these cells(Zhang et al. 2018b). We next asked if we can validate our single cell data and the absence of CAMK2A expressing cells in E-YTH but not in E-YTH^mut^ hippocami. While our single cell data did not reveal any CAMK2A expressing cells following *E- Yth* injections, Western blot analysis still detected a substantial, albeit reduced amount of CAMK2A proteins in E-YTH (Supplemental Fig. S4D). This discrepancy arises as our single cell experiments involved FACS sorting where we selected only successfully *E-Yth* transduced cells. In contrast, for Western blot, we examined the entire hippocampus due to material constraints, which includes many non-transduced cells. To further corroborate our findings, we also used an AAV virus (pAAV-*Camk2a*-*mCherry*) to visualize CAMK2A expressing cells, which appear red. This virus was mixed with equal amounts of EGFP expressing *E-Yth* or *E-Yth^mut^* AAVs and subsequently injected into mouse hippocampi. When we co-injected *E-Yth^mut^*and pAAV- *Camk2a*-*mCherry,* EGFP and mCherry signals overlapped, indicating the presence of CAMK2A expressing cells. However, when co-injecting *E-Yth* and pAAV-*Camk2a*-*mCherry,* the overlap was reduced, suggesting fewer or no CAMK2A expressing cells in our E-YTH single cell data and reinforces our single cell findings. Furthermore, the absence of CAMK2A expressing cells in E- YTH but not in E-YTH^mut^ suggests that m^6^A containing transcripts might be of particular importance in CAMK2A neurons (Supplemental Fig. S4E).

**Figure 3.**
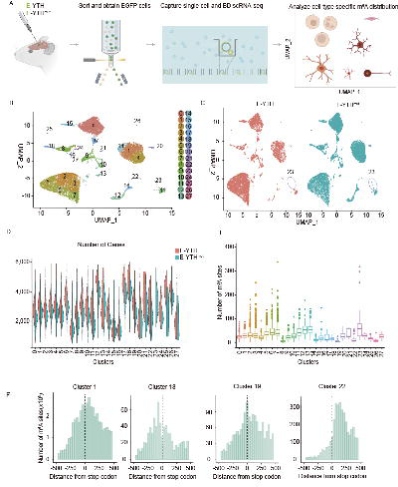
Hippocampal single cell identification following AAV transduction. (A) Schematic diagram of detecting m^6^A RNA modification in the mouse hippocampus on a single cell level through AAV transduction with *E-Yth* and *E-Yth^mut^* controls. (B) Integration of uniform manifold approximation and projection (UMAP) of 27,804 single cell transcriptomes. Cluster numbers from 0-27 are indicated. (C) Separate UMAP for E-YTH and E-YTH^mut^. 11,561 single cells in E-YTH and 16,243 in E-YTH^mut^. Circle: cluster 23 is missing in E-YTH. (D) Violin plot visualizing the number of genes per cluster. No genes were detected in cluster 23 in E-YTH samples. Orange: E-YTH; Blue: E-YTH^mut^. (E) Boxplot showing m^6^A counts per cluster. (F) Metagene analysis of individual cell clusters identified by single cell sequencing. m^6^A density surrounding the stop codon (position 0) is shown. m^6^A sites were obtained after eliminating background editing sites. Clusters 1, 18, 19 and 22 are shown that represent different m^6^A distribution patterns.

### Identification of m^6^A sites in single cell clusters of the hippocampus

Next, we identified C-to-U editing sites in our mouse hippocampus single cell sequencing data. To discover m^6^A sites in the single cell data, we first removed bulk background C-to-U editing events detected in mock, E-APOBEC1 and E-YTH^mut^ controls from our E-YTH single cell data (see Methods). Eventually, this left us with 2,566,141 C-to-U editing events that could represent m^6^A sites in our E-YTH single cell data. Despite removing this bulk background from our single cell E-YTH, due to unequal background distributions in clusters, it is also essential to remove cluster specific E-YTH^mut^ background detected in single cell E-YTH^mut^ (for more information see Methods). Thus, for each individual cell in our E-YTH single cell data, we subsequently removed the average cluster specific background that each cell belongs to. This provided us with 923,249 of m^6^A sites with transcripts on a single cell level (Supplemental Table S9). The highest number of m^6^A sites per cell, 178 (cluster average), was identified in cluster 22, while the smallest number of m^6^A sites per cell, 18 (cluster average), was identified in cluster 18 (Fig. 3E; Supplemental Table S9). To corroborate our single cell m^6^A data, we pooled our m^6^A single cell data and compared this with m^6^A sites identified by m^6^A-RIP and our bulk E-YTH data (Supplemental Fig. S3D). Although we found m^6^A overlaps across all methods, our single cell data has uncovered more m^6^A sites. This is not surprising, given its increased sensitivity.

We next analyzed the m^6^A distribution near the TTS. When merging all clusters together, we confirmed that the m^6^A distribution near the TTS was like bulk m^6^A sequencing data (Supplemental Fig. S5A). Although most m^6^A distribution patterns reflected what was observed with bulk m^6^A detection, we found cluster specific differences (Fig. 3F; Supplemental Fig. S5B). While cluster 1 had the most common m^6^A distribution pattern observed in most clusters, with the highest enrichment after the TTS, but also showed a smaller peak within 200bp before the TTS (Fig. 3F; Supplemental Fig. S5B). This pattern was reflected in the bulk m^6^A distribution pattern (Supplemental Fig. S5A). In contrast, a few clusters such as 22 and 5 lacked the smaller enrichment peak before the TTS, while in cluster 19 the m^6^A peak was at the TTS and in cluster 18 the m^6^A peak was upstream of the TTS (Fig. 3F; Supplemental Fig. S5B). Such distinct m^6^A distribution pattern, which are usually masked in bulk data, could be indicative of m^6^A regulatory or even m^6^A functional differences in specific cell types. This illustrates the advantages of detecting modifications at single cell resolution.

### Identification of 5 major cell types in hippocampal m^6^A single cell data

To determine which cell cluster corresponds to which cell type, we used automatic annotation (see Methods). We further verified our annotation criteria and gene markers with published work (Ximerakis et al. 2019). Overall, we determined the gene expression of cell type specific markers and grouped clusters that shared cell type specific marker expressions (for UMAP plots of all genes see Supplemental Material S1; Fig. 4A-C; Supplemental Table S10). Automatic annotation predicted cells of clusters 2,3,5,6,7,12 and 13 belong to oligodendrocytes, with cluster 16 being allocated to oligodendrocyte precursor cells. We verified that *Sox10*, a known marker gene for the oligodendrocyte cell (OLG) lineage, was indeed expressed in all these clusters(Ximerakis et al. 2019). Cells of cluster 22 were identified as astrocytes (ASC), where the astrocyte specific gene *Gja1* was exclusively expressed(Cid et al. 2021)(Fig. 4B). Cells of cluster 11 and 23 were identified as belonging to the neuronal cell lineage (NEU), as they expressed the neuronal cell lineage marker *Snap25* (Ximerakis et al. 2019). Cluster 26 contains endothelial cells (EC), expressing the marker gene *Esam* (Ximerakis et al. 2019). Clusters 0, 1, 4, 8-10, 14-15, 17- 21, 24-25, 27 were identified to belong to the immune cell (IMC) lineage (Ximerakis et al. 2019) (Fig. 4A-C). All cells in this group highly expressed *Lcp1*, but they could be subdivided into the following subcategories: (i) microglia, cluster 1, 4, 14, 20, with well-known marker gene expressions *Aif1*(*Iba1*), *Tmem119* and *P2ry12* (Jurga et al. 2020); (ii) Myeloid cells, cluster 0, 15, 17, 9, 10 and 27 (Ximerakis et al. 2019); (iii) B-cells, cluster 21 and 25, with high *Ly6d* expression (BLUMBERG 1990; Ximerakis et al. 2019); (iv) T-cells, cluster 18, 8, 24 and 19 with high *Cd3e* expression (BLUMBERG 1990).

**Figure 4.**
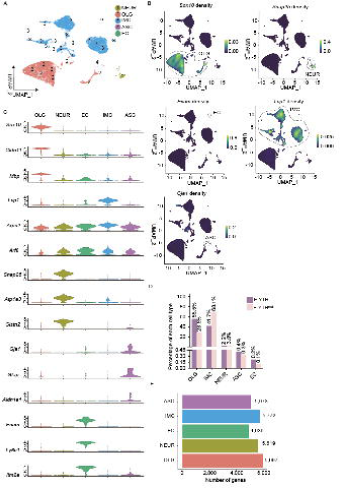
Hippocampal single cell type identification. (A) Uniform manifold approximation and projection (UMAP) with 5 main cell type populations were annotated and color coded based on cell type identifications. OLG: oligodendrocyte cell lineage; NEUR: neuronal cell lineage; EC: endothelial cell lineage; IMC: immune cell lineage; ASC: astrocyte cell lineage. (B) UMAP with expression levels of cell-type specific marker genes identifying all 5 major cell populations. Legend colour represents RNA density. Circles were added to visualize grouped cell populations. OLG: oligodendrocyte cell lineage cells have high expression of *Sox10;* NEUR: neuronal cell lineage cells have high expression of *Snap25;* EC: endothelial cell lineage cells have high expression of *Esam;* IMC: immune cell lineage cells have high expression of *Lcp1;* ASC: astrocyte cell lineage cells have high expression of *Gja1*. (C) Violin plot showing the distribution of expression levels of well-known representative cell-type-enriched marker genes across 5 cell types, 27,804 cells in total. (D) Histogram showing percentage of each cell type. (E) Histogram with number of detected genes per cell type.

In summary, our analysis led to the identification of 27,804 cells, which represent 5 main cell type lineages (Fig. 4A). We identified cells part of the oligodendrocyte (OLG) lineage, astrocytes (ASC), endothelial cells (EC), immune cells (IMC), as well as cells from the neuronal (NEUR) lineage. The most abundant cells identified in our single cell data are part of the OLG lineage, followed by IMC, NEUR, ASC and EC (Fig. 4D,E; Supplemental Fig. S6A; Supplemental Table S11). The IMC lineage contains notably more cells in E-YTH^mut^ controls versus E-YTH, although we did not notice any absent cell clusters.

### Hippocampal m^6^A cell type characteristics

Next, we intended to investigate cell type specific m^6^A characteristics. We identified 443 cells with an average of 7,667 m^6^A sites for ASC (178 sites per cell), 23 cells with an average of 595 m^6^A sites for EC (25 sites per cell), 487 cells with an average of 274,170 m^6^A sites for IMC (61 sites per cell), 232 cells with an average of 28,112 m^6^A sites for NEUR (121 sites per cell), 5,972 cells with an average of 612,705 m^6^A sites in OLG (102 sites per cell) (Fig. 5A,B; Supplemental Fig. S6B; Supplemental Table S12). Next, we generated UMAP plots for every gene where we detected m^6^A sites, illustrating the total number of m^6^A sites on RNA transcribed from a specific gene (Supplemental Material S2). Based on the UMAP plots and depending on the gene, we noticed that many m^6^A sites seem to be differentially expressed (Supplemental Material S2; Fig. 5C). To have confidence in the differential m^6^A patterns in our single cell data, we initially intended to test if we can verify these single cell findings other ways. To do this, we focused on two abundant cell types for ease of isolation, IMC and OLG (see Methods). Using our single cell data, we identified *Colgalt1* and *Gsn* as genes expressing differentially methylated transcripts in IMC and OLG (Fig. 5C). After isolating the relevant cell groups form wild type mouse brain hippocampi, we performed m^6^A-RIP, followed by reverse transcription and qPCR. This process allowed us to quantify the enrichment of *Colgalt1* and *Gsn*, representing their m^6^A abundance in IMC and OLG cells (Fig. 5D). Our m^6^A-RIP demonstrates that *Colgalt1* transcripts have detectable m^6^A sites in IMC, but not in OLG. Conversely, *Gsn* showed a higher enrichment for m^6^A sites in IMC than in OLG (Fig. 5E). This m^6^A-RIP experiment substantiates our single cell m^6^A findings and enhances the reliability of our conclusions (Fig. 5C-E).

**Figure 5.**
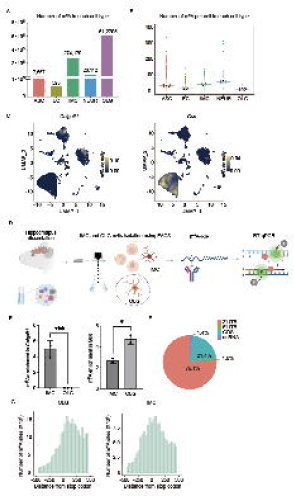
m^6^A single cell distribution per hippocampal cell type. (A) Boxplot showing m^6^A counts per cell type. ASC: astrocyte cell lineage. EC: endothelial cell lineage. IMC: immune cell lineage. NEUR: neuronal cell lineage. OLG: oligodendrocyte cell lineage. (B) Boxplot with m^6^A counts per cell for each cell type. Number in box plot reflects average. (C) UMAP illustrating the m^6^A density on RNA transcribed from one gene, per cell. Plots for genes *Colgalt1* and *Gsn* are shown. Legend colour represents density of m^6^A on RNA transcribed. (D) Schematic diagram of approach to confirm differential m^6^A RNA modifications in different cell populations. Relevant cell groups form wild type mouse brain hippocampi are isolated by FACS sorting, such as IMC versus OLG, followed by m^6^A RNA immunoprecipitation (m^6^A-RIP), reverse transcription and qPCR (RT-qPCR). m^6^A-RIP enrichment representing relative m^6^A abundance in in different cell populations. OLG: oligodendrocyte cell lineage. IMC: immune cell lineage. (E) Transcript m^6^A enrichment quantifications in IMC and OLG populations following m^6^A-RIP and RT-qPCR versus input control samples. Transcript m^6^A enrichments represent m^6^A abundance in IMC and OLG cells. Both transcripts of *Colgalt1* and *Gsn* were detected in all input control samples. No *Colgalt1* transcripts were detected following m^6^A-RIP in OLG. OLG: oligodendrocyte cell lineage; IMC: immune cell lineage. (F) Pie chart showing m^6^A distribution within RNA. Data represents pooled single cell data. CDS: coding site. ncRNA: non-coding RNA. (G) Distribution of m^6^A surrounding the stop codon (0nt) identified by single cell sequencing for OLG and IMC. OLG: oligodendrocyte cell lineage; IMC: immune cell lineage.

To determine if the m^6^A differential distribution could be explained by different levels of m^6^A regulatory enzymes, we also analyzed the level of some m^6^A regulatory enzymes at single cell level (Supplemental Fig. S6B). Although we found association between some m^6^A methylases and m^6^A increased density, we also saw a higher level of the m^6^A demethylase *Alkbh5* (Supplemental Fig. S6B). This suggests that m^6^A regulation cannot be explained by the main regulator enzymes alone. This indicates that m^6^A regulation is more complex than currently understood, which is tightly controlled, in a cell specific manner. Next, we analyzed our single cell data further to discover more cell type specific differences. Although our pooled single cell data confirmed previously reported m^6^A mRNA distribution patterns(Dominissini et al. 2012; Meyer et al. 2012), it does not reveal novel insights (Fig. 5F; Supplemental Table S13). Thus, we focused on the cell type level.

OLG, as well as NEUR, ASC and EC showed a major m^6^A peak shortly after the TTS (Fig. 5G; Supplemental Fig. S6C). This pattern was also reflected in the bulk m^6^A distribution pattern (Supplemental Fig. S5A). In contrast, IMC cells also showed a smaller m^6^A peak within 200bp before the TTS, in addition to the high m^6^A enrichment after the TTS (Fig. 5G; Supplemental Fig. S6C). Such distinct m^6^A patters could be indicative of m^6^A regulatory or even m^6^A functional differences between cell types.

### Heterogeneous and homogeneous m^6^A sites in single cells

We next asked how often m^6^A occurs at a particular site on RNA replicates in our single cell data, reflected by the m/k ratio per gene per cell. A high m/k ratio is indicative of homogenous m^6^A sites, while a low m/k ratio suggest RNA replicates have few m^6^A per gene per cell. Upon investigating potential variations in m^6^A density m/k distributions between clusters and cell types, we found strikingly similar distributions across all categories (Fig. 6A; Supplemental Material S3). Despite the presence of heterogenous m^6^A sites (m/k<0.25), we identified a greater number of homogenous m^6^A sites, with an m/k=1. To verify our m^6^A homogeneity findings, we compared our single cell m^6^A m/k distribution with single cell E-YTH^mut^ controls, which showed much more heterogeneity and fewer homogenous (m/k=1) m^6^A sites (Fig. 6A; Supplemental Material S3). Like our bulk data, we have removed background noise, such as small nuclear polymorphisms (SNPs) and E-YTH^mut^ background. Additionally, here, we wanted to ensure that the homogenous (m/k=1) sites we observed were not mouse-specific SNPs. Thus, we identified all m/k=1 m^6^A sites per cell types, and then only kept those sites that occurred in other cells and with a lower m/k ratio (m/k<0.9). Although this decreased our homogenous m^6^A gene list, we ensured that our homogenous m^6^A were not SNPs (Supplemental Table S14). We next evaluated these conserved homogenous (m/k=1) m^6^A sites (Fig. 6). We asked if m/k=1 m^6^A sites are differentially distributed than more heterogenous (m/k<1) m^6^A sites. Indeed, we found that m/k=1 and m/k<1 sites show clear distribution differences between each other in OLG and IMC groups. In OLG, while heterogenous m^6^A are mostly enriched shortly after TTS, most homogenous m^6^A sites are enriched just before the TTS, with another m^6^A peak at the TTS site itself (Fig. 6B). In contrast, heterogenous IMC m^6^A sites are enriched after the TTS, illustrated by one major m^6^A enrichment peak. Although homogenous IMC m^6^A sites also occur after the TTS, they are distributed more evenly along the 3’UTR (Fig. 6B). These different distribution patterns could be indicative of cell type specific m^6^A regulation and potentially function. To obtain an insight into the kind of transcripts and pathways that could be regulated by homogenous m^6^A sites, we performed gene ontology analysis (Fig. 6C; Supplemental Fig. S7A). We identified many lineage-specific pathways, such as pathways affecting axons for NEUR, oligodendrocyte specific pathways for OLG or postsynaptic pathways for ASC (Supplemental Fig. S7A). We also identified disease pathways such as Amyloidosis, Dementia and Degeneration for NEUR, or Hydrocephalus and Amyloid Plaque for ASC (Fig. 6C; Supplemental Fig. S7B). We next focused on the genes in which m^6^A occurs with an m/k=1 in at least once cell (Supplemental Table S14). First, we noticed that m^6^A occurs on transcripts coding for its m^6^A demethylase ALKBH5, suggesting a regulatory feedback mechanism (Supplemental Table S14). Second, we identified that m^6^A is also present in both *Fos* and *Jun* in OLG and IMC, and that m^6^A can also occur at multiple sites on one transcript, such as on *Stat1* (Supplemental Table S14). Noticeably, we found that *Smarcc2* RNA also has m^6^A sites (Supplemental Fig. S8; Supplemental Table S14). SMARCC2, is a member of the SWI/SNF family of proteins that regulate chromatin structure. It is known to be implicated in neural stem cell proliferation and in the transition between proliferating neural progenitor cells to postmitotic neurons (Nguyen et al. 2018). Our finding suggests that m^6^A might be implicated in this regulatory mechanism through SMARCC2. Strikingly, we also identify several genes that are associated with Alzheimer Disease (AD) to have many cells with an m/k=1 ratio, such as *App*, but also on *Apoe*, *Aplp1*, *Ctsb* and *Itm2b* (Fig. 6D; Supplemental Fig. S8; Supplemental Table S14)(Turner et al. 2003; Priller et al. 2006; Hook et al. 2023). Our findings suggest that *App*, *Apoe*, *Aplp1*, *Ctsb* and *Itm2b* might be regulated by m^6^A. We also found other m^6^A transcripts with m/k=1 to be associated in diseases, such as *Mecp2* (encoding the methyl CpG binding protein 2) in NEU, Synaptotagmin-11 (*Syt11*), *Lamp1* and *Brd2* (Supplemental Fig. S8; Supplemental Table S14). Mutations in *Mecp2* gene cause the neuronal disease Rett Syndrome (Guy et al. 2007), *Syt11* is associated in Schizophrenia and Parkinson Disease (Inoue et al. 2007; Lill et al. 2012), LAMP1 is a Lassa virus receptor (Enriquez et al. 2022), while *Brd2* is associated with epilepsy (Pal et al. 2003) (Supplemental Fig. S8; Supplemental Table S14). Overall, these identified transcripts with m^6^A sites that occur in all RNA replicates in a specific cell are likely to be particularly sensitive to m^6^A regulation.

**Figure 6.**
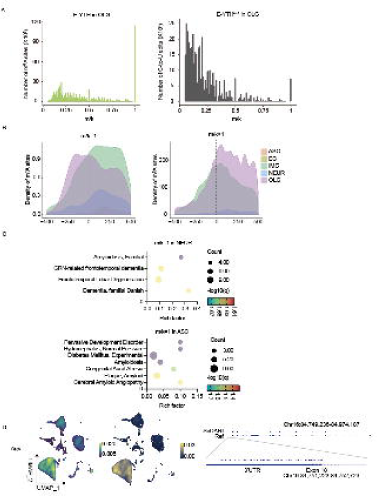
Heterogeneous and homogeneous m^6^A sites in single cells. (A) Histogram of m^6^A site counts over mutation per read (m/k) ratio and histogram of C-to-U edit E-YTH^mut^ background over mutation per read (m/k) ratio. The data represents single cell data for OLG. OLG: oligodendrocyte cell lineage. A minimum threshold of 5% was applied. (B) Metagene analysis showing normalized m^6^A scaled density 500nt 5’ and 500nt 3’ from stop codon (0nt) for ASC, EC, IMC, NEUR and OLG. Left: m^6^A sites with m/k=1 (homogenous m^6^A). Right: m^6^A sites with m/k<1 (heterogenous m^6^A). m/k: mutation per read ratio. OLG: oligodendrocyte cell lineage; NEUR: neuronal cell lineage; EC: endothelial cell lineage; IMC: immune cell lineage; ASC: astrocyte cell lineage. (C) Disease gene enrichment analyses (GO). GO terms with an adjusted q-value<0.05 and p-value<0.05 are plotted for NEUR and ASC. NEUR: neuronal cell lineage; ASC: astrocyte cell lineage. (D) Localization of m^6^A for gene *App* that is associated with Alzheimer Disease is shown. Left: UMAP plot of single cell expression for one gene. Legend colour represents transcript density. Middle: UMAP density plot of m^6^A on RNA transcribed from one gene, per cell. Legend colour represents m^6^A density on RNA transcribed from one gene. Right: IGV RefSeq gene annotations with editing sites representing adjacent m^6^A sites is shown. Last exon with 3’UTR region is illustrated with higher magnitude.

In conclusion, our single cell data of the hippocampus identified an array of different transcripts and cell clusters and cell types that are likely to be regulated by m^6^A. Our landscape of m^6^A modifications within individual cells provides new insights into the molecular mechanisms underlying hippocampal physiology and lays the foundation for future studies investigating the dynamic nature of m^6^A RNA methylation in the brain.

## DISCUSSION

In this study, we optimized the DART-seq system (Meyer 2019) to identify m^6^A sites in single cell data from the mouse hippocampus. Initially, we generated *E-Yth* and *Yth-E* constructs and their respective controls, testing their efficacy in detecting m^6^A sites in RNA from HEK293T cells. Our findings align with previous m^6^A studies, which were also validated through functional investigations(Meyer 2019) We determined that the *E-Yth* construct exhibited superior performance, showing improved detection of *E-Yth* transfected cells and stronger correlation between EGFP signal and APOBEC1-YTH presence. By using the *E-Yth* construct, we achieved a more distinct m^6^A distribution pattern by excluding cells with low E-YTH expression, thereby reducing background noise. Subsequently, we packaged the *E-Yth* construct and its corresponding *E-Yth^mut^* control into AAVs and injected them into mouse hippocampi. We isolated transduced cells and performed bulk and single cell sequencing analysis to identify m^6^A sites.

Overall, we have identified 923,249 m^6^A sites within transcripts with single cell resolution in the mouse brain hippocampus. We found CAMK2A expressing cells missing after APOBEC1- YTH treatment. It seems APOEBC1-YTH interferes with essential m^6^A transcripts in CAMK2A expressing neurons, suggesting that m^6^A containing transcripts might be of particular importance for CAMK2A neurons. Alternatively, APOBEC1-YTH could cause other abnormalities. Since APOBEC1-YTH causes C-to-U edits in the vicinity of m^6^A sites facilitating their identification (Fig. 1A), as such, it also introduces mutations in the RNA. Alternatively, APOBEC1-YTH binding to m^6^A sites itself, but not the mutation it introduces, could interfere with the function of specific m^6^A transcripts or compete with m^6^A reader proteins such as YTHDF2, leading to changes in phenotypes. Although C-to-U mutations nearby m^6^A targets have been previously reported not to interfere with transcriptional changes or cell viability (Meyer 2019), occasionally, such mutations could result in functional changes or even cell death, which might explain the E-YTH specific absence of CAMK2A neurons. In these cells, the E-YTH seems to interfere with the transcriptome in such a way that either the transcriptome can no longer be allocated to CAMK2A neurons, or that the cells are missing due to decreased cell viability. Overall, it appears that in CAMK2A neurons, m^6^A or the transcripts that contain m^6^A are of particular importance.

The CAMK2A protein is found at the synapse. It regulates synaptic transmission, excitability and long-term potentiation in hippocampal neurons, which play an important role in learning and memory (Yasuda et al. 2022). CAMK2A and CAMK2B are two of the most abundant proteins in the Post Synaptic Density (PSD) regions within the hippocampus, making up approximately 10% of total protein in the PSD(Yasuda et al. 2022). Interestingly, METTL3 depletion only in CAMK2A expressing cells in the mouse hippocampus was shown to reduce long-term memory consolidation, highlighting the functional and biological importance of these cells(Zhang et al. 2018b). Also, reports have shown depression-like behavior, decreased spatial memory altered fear conditioning when the METTL3 m^6^A methylase is depleted in mice(Engel et al. 2018; Xu et al. 2022). Since these phenotypes are as well regulated by the hippocampus and by CAMK2A neurons, the absence of CAMK2A cells in our E-YTH single cell data but not in our E-YTH^mut^ controls corroborates that the phenotypes associated with m^6^A deregulation are indeed due to abnormalities in m^6^A transcripts of CAMK2A expressing cells.

Our study also reveals that although on a population level m^6^A is heterogenous, there are many m^6^A sites that are homogenous within a cell. This is surprising, as previous reports have reported single cell level m^6^A heterogeneity in HEK293T cells (Tegowski et al. 2022) and our bulk data also showed similar results. In contrast, the single cell data revealed homogeneity of m^6^A sites and identified differential distribution patterns near TTS between homogenous and heterogenous m^6^A in some cell lineages. This could be indicative of regulatory and functional differences. We hypothesize, that m^6^A RNA targets that have low m/k values, their function might overall be only fine-tuned by m^6^A regulators and cause no or limited phenotypic changes. In contrast, transcripts with high m/k ratios, such as *Smarcc2,* might be particularly sensitive to m^6^A regulation. Here, we also discover many such m^6^A transcripts in specific cell types that are associated with brain diseases, such as Alzheimer Disease (AD), Schizophrenia, Rett Syndrome or Epilepsy. An earlier study reported a decrease in m^6^A levels with age and in AD patients found a link between m^6^A levels and transcripts linked to synaptic function such as *Camk2* (Castro-Hernández et al. 2023) This, together with our findings suggests that some specific m^6^A transcripts such as *App*, *Apoe*, *Aplp1*, *Ctsb* and *Itm2b* and m^6^A transcripts in CAMK2A neurons of the hippocampus might not only sensitive to m^6^A regulation, but are likely to be implicated in ageing and AD pathology.

By investigating m^6^A RNA methylation on a single cell level, we uncover novel insights into the hippocampus and provide a deeper understanding of the m^6^A transcriptome in this critical brain tissue. As we identify cell-type and transcript specific m^6^A sites, these sites could be used as future potential therapeutic targets. Our work lays the foundation for future studies investigating the dynamic nature of m^6^A RNA methylation in the healthy and diseased brain.

## METHODS

### Cell lines

HEK293T cells were purchased from Beijing Xiehe Cell Bank and cultured at 37°C with 5% CO2 in DMEM (Dulbecco’s Modified Eagle Medium) containing 10% FBS and 1% penicillin/streptomycin. All experiments evaluating any AAV vectors were performed in the HEK293T cell line. Cells were passaged for less than 20 times and have been regularly tested for mycoplasma.

### Vector cloning and Adeno-associated virus

An Adeno-associated plasmid pAAV-*Cag-Egfp-Wpre-Sv40* (gift from Minmin Luo) was used as the backbone to generate the viral expression constructs pAAV-*Cag-Apobec1-Yth-Egfp* (*Yth-E*, Addgene #209322), pAAV-*Cag*-*Apobec1*-*Yth^mut^*-*Egfp* (*Yth^mut-^E*, Addgene #209323), pAAV-*Cag-Apobec1*-*Egfp* (Addgene #209324) pAAV-*Cag*-*Egfp*-*Apobec1*-*Yth* (*E-Yth*, Addgene #209319), pAAV-*Cag-Egfp-Apobec1*-*Yth^mut^*(*E*-*Yth^mut^*, Addgene #209320) and pAAV*-Cag-Egfp-Apobec1* (Addgene #209321). To clone these vectors, the cassette pCMV-*Apobec1-Yth* (Addgene #131636) and pCMV-*Apobec1-Yth^mut^* (Addgene #131637) were inserted into the backbone of pAAV-*Cag-gfp*-*Wpre-Sv40* using In-Fusion cloning. We also used pAAV-*Camk2a*- *mCherry* (gift from Fei Zhao). In brief, the plasmid to be packaged was co-transfected into HEK293T cells with a rep/cap containing plasmid pUCmini-iCAP-PHP.eB (Addgene #103005) and the helper plasmid pAdDeltaF6 (Addgene #112867), in the presence of Polyethyenimine. After 72hrs the produced AAV virus was harvested, purified by Chloroform, titrated and quantified by qPCR (Negrini et al. 2020a). AAV stereotaxic injections were performed targeting the hippocampus of 3-month-old mice. The following positions were used: X/P: 1.94mm, M/L: 1.5mm from the bregma point with a depth of 2mm.

### Immunostaining and Microscopy

HEK293T cells were seeded on 35mm-diameter dish (WPI’s FluoroDish™), 5µg plasmids were transfected and images were taken after 24 hours. Cells were fixed with 4% paraformaldehyde for 10 mins and wash twice with PBS followed by 15min permeabilization with 0.1% triton-x100. Following PBS washes, the samples were blocked for 1 hour at RT in PBS with 1% BSA, incubation at 4 °C overnight with the HA monoclonal antibody (Alexa Fluor™ 555, Invitrogen # 26183-A555). After 3 PBS washes and 0.1% DAPI staining, fluorescent images were captured using Zeiss Inverted confocal microscope, using the ZEN blue software (DAPI: Excitation 365, BS FT 395, Emission BP 445/50; GFP: Excitation BP 470/40, BS FT 495, Emission BP 525/50; CY3: Excitation BP 545/25, BS FT 570, Emission BP 605/70).

Mouse hippocampi slices in Fig. 2B were imaged with the Olympus VS120 Virtual Slide Microscope, using the OlyVIA software (DAPI: Excitation 365/10 nm, Emission 440/40 nm; GFP: Excitation 472/30 nm, Emission 520/35 nm). Mouse hippocampi slices in Supplemental Fig. S4 were captured using the Olympus VS200 Virtual Slide Microscope, using the OlyVIA software (Cy3: Excitation 555/20 nm, Emission 595/33 nm; GFP: Excitation 480/30 nm; Emission 519/26 nm; DAPI: Excitation 395/25 nm, Emission 434/32 nm).

### Bulk m^6^A sequencing in HEK293T cells

HEK293T cells were cultured and processed independently of each other to generate independent biological replicates. Three independent replicates were processed for *Yth-E* and *Yth^mut^-E* expressing cells. 24 h after plasmid transfections, cells were rinsed with DPBS and digested using 0.5% Trypsin-EDTA. Cells were pelleted, washed with DPBS and filtered through a 40-µm cell filter prior to FACS analysis. Cells were loaded onto a custom FACS ARIA III flow sorter (BD Biosciences) equipped with a forward scatter photomultiplier tube. Single gating was used to isolate single cells. Intact cells were selected by gating the EGFP signal. The EGFP signal was detected by green excitation light (488nm laser, green FITC). The nozzle size of the flow cytometer was 100µm. Particles smaller than cells (black dots) were eliminated using the forward scatter (FSC-PMT-A) *versus* side scatter (SSC-A). Cell-sized particles were gated (box). Plots of height *versus* width in the forward scatter and side scatter channels were used to exclude aggregates of two or more cells. Following FACS isolation of EGFP positive single cells, total RNA was isolated with the Micro Total RNA Isolation Kit (Invitrogen, #AM1931) according to the manufacturer’s instructions. Total RNA was treated with DNase I (Tiangen, RT411) for 15 mins at room temperature to remove DNA contamination. Sequencing libraries were generated from 1-10ng of total RNA from each replicate using the Single Cell Full Length mRNA kit (Vazyme, #N712) and TruePrep® DNA Library Prep Kit V2 (Vazyme, #TD502), according to the manufacturer’s instructions. Quality control was performed using the Fragment Analyzer Systems Capillary Arrays (Agilent, FA12) and quantified with Qubit 1X dsDNA HS Kit (Invitrogen, #Q33231). 150bp paired-end sequencing was performed on a NovaSeq 6000 (Illlumina) using a S4 flow cell.

### Identification of m^6^A sites in bulk RNA sequencing

Low-quality bases (<Q20) and adaptor sequences were trimmed using Trimmomatic (0.39) (Bolger et al. 2014), and reads with less than 36 nucleotides were subsequently discarded. The remaining reads were aligned to the mm39 reference genome using bwa-mem (0.7.17) (Li and Durbin 2009). Duplicate reads were marked using the MarkDuplicates tool from Picard (see Method). Subsequently, CLIP Tool Kit (CTK) was utilized to collapse PCR duplicates and to identify C-to-U editing events, following default parameters (Shah et al. 2017).

To identify C-to-U and m^6^A sites, we used an approach as described by Meyer *et al*. (Meyer 2019), using the following filters: (i) Only C-to-T mutations with a false discovery rate (FDR) of less than 0.01 were kept for any downstream analyses; (ii) Only C-to-T mutations with two or more editing events (m ≥ 2) and a coverage of at least 10 (k ≥ 10) was considered for downstream analyses; (iii) C-to-T mutations with a ratio of m/k (Number of reads with a C-to-T mutation / total reads per site) greater than 5% were kept for downstream analyses; (iv) C-to-T mutations sites that were found in single nucleotide polymorphisms (SNPs) databases Mouse Genome Project (mgp_REL2021_snps) and Genome Reference Consortium Mouse Build 39 (GCA_000001635.9) were discarded; (v) Only C-to-T mutation sites that were found in at least two out of three replicates were considered for downstream analyses.

To identify m^6^A sites in the C-to-T converted YTH-APOBEC1 sequencing data, we first removed any C-to-T conversions sites that were also identified in our wild type (mock) and APOBEC1 overexpression background control samples. Second, we only kept C-to-T conversions sites in the YTH-APOBEC1 sequencing data that occurred at least ≥1.5 times more frequent than in the YTH^mut^-APOBEC1 negative control sequencing data. Overall, all the above restrictions allowed us to remove false positives and leave us with stringent C-to-T conversion sites in our YTH-APOBEC1 datasets. These remaining C-to-T conversion sites in the YTH-APOBEC1 data represent our m^6^A sites, which we refer to as m^6^A sites in the bulk data.

### Analyses and plotting

Biorender was used for some illustrations, as well as IGV and Prism. All experiments were carried out with 3 technical and biological replicates, indicated by n. Statistical analyses and plots were performed using R (ref). The VennDiagram package was used for Venn diagrams, Seurat for UMAP plots and ggplot2 for the rest of the plots. metaPlotR was used to generate m^6^A metagene plots, such as histograms along simplified transcript models, of the C-to-U conversion (Olarerin-George and Jaffrey 2017). When multiple transcript isoforms could potentially contain the C-to-U site, the longest isoform was chosen.

### Mass spectrometry

Analysis of global levels of A and m^6^A was performed on a TSQ Altis^TM^ triple quadrupole mass spectrometer (Thermo Fisher Scientific) coupled to a Vanquish Flex UHPLC system (Thermo Fisher Scientific) fitted with an Acquity UPLC HSS T3 column (2.1 x 100 mm, 1.8 μm particle size, Waters). The mobile phase consisted of 0.5% aqueous formic acid (solvent A) and 0.5% formic acid in acetonitrile (solvent B) at a flow rate of 300 µl/min. Calibration curves were generated using serial dilutions of synthetic standards for adenosine (A, Sigma-Aldrich) and N6-methyl-2’-adenosine (m^6^A, Sellechchem). The mass spectrometer was set in a positive ion mode and operated in selective reaction monitoring. The precursor ions of A (m/z 268.1) and m^6^A (m/z 282.1) were fragmented and the product ions of A (m/z 136.1) and m^6^A (m/z 150.1) were monitored. The EIC of the base fragment was used for quantification. Accurate mass of the corresponding base-fragment was extracted using the XCalibur Qual Browser and XCalibur Quan Browser software (Thermo Fisher Scientific) and used for quantification. Quantification was performed by comparison with the corresponding standard curve obtained from the pure nucleoside standards running with the same batch of samples. The level of m^6^A present in the sample was expressed as a percentage of total adenosine content (methylated and non-methylated), calculated according to the following equation: (%) m^6^A = 100 x m^6^A /[A]. Differences in m^6^A percent abundance were considered significant when P ≤ 0.05.

### Mouse strains

Animals were maintained and processed in accordance with CIBR guidelines. All experimental methods were approved and followed all regulations of the Welfare and Ethics Review Committee for Laboratory Animals. C57BL/6J mice were originally purchase from the Jackson Laboratory but were made available through CIBR’s animal facility. All mice were housed in a 12:12 light-dark cycle, under controlled climate and enrichment environmental conditions with access to sterile food and water *ad libitum*.

### m^6^A RNA immunoprecipitation

Total RNA was extracted from the hippocampus of adult mice (C57BL/6J) using Trizol (Thermo-Fisher) reagent. After removing genomic DNA, Qiagen Rneasy kit was used for RNA purification, resulting in around 20μg of total RNA per mouse. The integrity of RNA was assessed using the Fragment Analyzer Systems Capillary Arrays (Agilent F12), while the concentration and purity were determined using a spectrophotometer (Thermo Nanodrop one). For RNA fragmentation, the samples were incubated for 4 minutes at 94°C in a fragmentation buffer (containing 10mM ZnCl2, 10mM Tris-HCl, pH 7), followed by standard isopropanol precipitation. For m^6^A RNA immunoprecipitation (RIP), an existing protocol was adjusted (Dominissini et al. 2012). ImmuProtein A Dynabeads (Invitrogen, #10001D) were washed three times with IP buffer (containing 150mM NaCl, 0.1% NP-40, 100mM Tris-HCl, pH 7.4 and 6 µg/µL BSA) and incubated with rotation in IP buffer for 2 hours at 4°C. Each RNA sample was divided to obtain an input control sample (10%) and the 90% was incubated with an anti-m^6^A antibody (SySy, #202011, 2.5μg). The RNA-antibody mixture was incubated for 2 hours at 4°C with rotation. The magnetic beads were then conjugated with the antibody-RNA solution, which selectively captures RNA fragments containing m^6^A modifications. After three washes with IP buffer, the antibody captured RNA was eluted for 1 hour at 4°C with rotation using elution buffer (containing IP buffer and 6.6mM m^6^A) and was then concentrated by isopropanol precipitation. To generate RNA-seq libraries for input control and m^6^A-RIP pulldown samples, the RNA was processed using the SMARTer Stranded Total RNA-Seq Kit v2 (Takara, #634411). The fragment length of the libraries was verified using the Fragment Analyzer 12 (Agilent). Paired-end 150- base pair reads were sequenced with the NovaSeq 6000 platform (Illumina).

### m^6^A RNA immunoprecipitation sequencing data analysis

rRNAs were removed using the mouse rRNA reference (GCF_000001635.27_GRCm39_rna_from_genomic.fna) from NCBI. Adapters were eliminated with Cutadapt (version 2.8)(Martin 2011). Additionally, the first ten 5’ nucleotides were eliminated from the ends due to the presence of potentially low-quality nucleotides. PCR duplicates were removed from the aligned datasets. Mapping and alignment was done by Hisat2 (2.2.1) (Kim et al. 2019), followed by peak calling using Macs2 (version 2.2.6) (Gaspar 2018).

### Hippocampi dissociation

We combined and adapted existing protocols to enable FACS sorting of intact single cells with high viability. Briefly, two weeks after performing AAV brain stereotaxic injections into mice hippocampi, mice were anesthetized and then perfused with Dulbecco’s phosphate buffer saline. Brains were dissected and hippocampi were extracted in cold DPBS solution containing calcium, magnesium, glucose. The hippocampi were dissociated into single cells using the Adult Brain Dissociation Kit (Miltenyi Biotex, #130-107-677), with the following optimized conditions: (i) approximately 250mg of adult hippocampi was used and was treated as one sample as input material; (ii) a gentle MACS program of >100mg:37°C_ABDK_01 was chosen; (iii) debris were removed following the manufacturer’s manual; (iv) 10ml PB buffer was used to suspend the cells with cold 1x Red Blood Cell Removal Solution; (v) PB buffer was used for FACS sorting.

### Single cell m^6^A sequencing

8 hippocampi from 3-month-old mice were pooled for each sample to obtain sufficient cell numbers per sample. E-YTH and E-YTH^mut^ EGFP positive cells were isolated through FACS sorting and processed into cDNA libraries. Briefly, after hippocampi dissociation, cells were FACS sorted and EGFP positive were collected in ice-cold PBS containing 0.5% BSA. Per sample, 30,000 cells were loaded into a Rhapsody Cartridge (BD Biosciences) and processed following the manufacturer’s instructions. Subsequently, single cell RNA sequencing libraries were prepared using the Rhapsody WTA kit (BD Biosciences). The Rhapsody, BD single cell platform was chosen for single cell processing as it seems gentler than the 10x platform. Libraries were pooled and then sequenced by NovaSeq 6000 (Illumina). The BD^TM^ Rhapsody docker setup (BD) was used to perform barcode processing and single cell gene-UMI counting, following manufacturer’s instructions (BD Rhapsody Sequence Analysis Setup v1.0). A digital expression matrix was obtained for each experiment with default parameters and was mapped to the mouse reference genome mm39.

### Single cell sequencing gene expression analysis

The reads in the fastq files from E-YTH and E-YTH^mut^ samples were aligned to the mm39 reference genome following BD Rhapsody^TM^ Sequence Analysis Pipeline v1.0. Matrices containing RSEC-corrected molecules per bioproduct per cell numbers were loaded into Seurat (3.1.5) (Butler et al. 2018). Low-quality cells with fewer than 500 or more than 6,000 detected genes were excluded. Additionally, cells with a high mitochondrial content (>10%), indicative of poor cell quality, were also filtered out. The remaining cells were considered for downstream analysis. The data from each individual sample was then log-normalized. The top 2,000 more variable genes within each sample were identified using the FindVariableFeatures function in Seurat and were used as integration anchors for the integration of E-YTH and E- YTH^mut^ samples. This was done using the FindIntegrationAnchors and IntegrateData functions in Seurat. Integrated data was subsequently scaled, and Principal Component Analysis (PCA) was performed to reduce the dimensionality of the data. The top 30 principal components were then used to identify cell populations or clusters across the two samples, using the FindNeighbors and FindClusters functions. The number of principal components to use was determined by analyzing the elbow plot. The plot revealed that after 30 principal components, the explained variance by each component started to level off. The resolution parameter in FindClusters function was set to 0.8. Uniform Manifold Approximation and Projection (UMAP) was subsequently applied to visualize the clustered cells in a two-dimensional space.

To determine cluster-specific marker genes, the FindConservedMarkers function in Seurat was employed, comparing each cluster against all other clusters within the integrated dataset. For each run of this function, only genes detected in at least 10% of the cells in either of the two populations were considered for analysis. Genes with a log-fold change greater than 0.3 were considered as cluster-specific markers. To help annotate the identified clusters, we followed two automatic cell-type annotation approaches. First, we used the scMCA function in the scMCA package (0.2.0) (Sun et al. 2019). This function assigns mouse cell types to cell clusters based on expression profiles. Second, we used SciBet (1.0) cell type classifier with the Tabula Muris Brain Non-Myeloid model (Li et al. 2020). Seurat was used for UMAP plots(Hao et al. 2021), and ggplot2 for the rest of plots(Wickham 2016). UMAP plots were generated with Nebulosa, using the plot_density function (Alquicira-Hernandez and Powell 2021).

### Differential gene expression analysis

Gene-level read counts were obtained from the aligned files using featureCounts (2.0.0) (Liao et al. 2014). Differential expression analysis between E-YTH and E-YTH^mut^ cells was performed using DESeq2 (Love et al. 2014). After filtering out genes with low expression levels (number of reads <10) and adjusted P-values >0.1, volcano plots were plotted. Adjusted P-values <0.05 and a Log2FC>1 and Log2FC<-1 are considered significant. Genes were identified as differentially expressed between the two conditions with fold change of Log2FC>1 and Log2FC<-1. (Adjusted P-value <0.05).

### Western Blot

Hippocampi were homogenized in 500µl of RIPA medium lysis buffer (Beyotime, #P0013E-2) in the presence of protease inhibitor cocktail (Roche, #11836170001). Samples were centrifuged at 10,000 × g for 30 minutes. Proteins were quantified with the Qubit Protein Assay Kit (Invitrogen, Q33212), dilute to 1 µg/µl followed by denaturation using 5 X SDS loading buffer (CWBIO, #CW0027) at 95 °C for 10 minutes. 10 µg of proteins were run on 10% PAGE gels (Vazyme, #E303-01) at 170V for 60 minutes. Proteins were transferred onto activated PVDF transfer membranes (Immobilon, #IPVH00010). Membranes were washed with TBST (10mM Tris-HCl, pH 8.0, 150mM NaCl, 0.05% Tween 20) once for 5 min at room temperature, followed by blocking 5% nonfat dried milk for 1 hour at room temperature. The membrane was incubated at 4°C overnight with antibodies such as CAMK2A (Thermo Fisher Scientific, #MA1-048) and GAPDH (Abcam, #ab9485), which were diluted in TBST with 5% BSA. After 3 washes with TBST, the membranes were incubated with HRP-conjugated secondary antibodies (Abcam, Ab205718 & Ab205719) in TBST with 5% BSA, for 1 hour at room temperature. After 3 washes the signals were visualized with an HRP substrate Peroxide solution and Luminol reagent (Immobilon #WBKL5S0500). iBright 1500 (Invitrogen) was used to detect and quantify protein presence. Background signal was removed (background corrected volume, Local Bg. Corr. Vol.) to quantify the relative amounts of proteins. CAMK2A levels were normalized against GAPDH for each biological replicate.

### Identification of m^6^A sites in single cell sequencing data

The software Bullseye was used to identify m^6^A sites in single cell sequencing data (Tegowski et al. 2022). We applied the same conditions for single cell analysis as we did for bulk data when identifying m^6^A sites, except for the following adjustments: We removed any C-to-U editing sites that were identified in the YTH-APOBEC1 and wild type bulk sequencing analysis from the YTH-E single cell data. We then selected only those that show a 1.5-fold increase over bulk E-YTH^mut^ controls. Since C-to-U background edits are not equally distributed amongst all clusters, it is essential to remove E-YTH^mut^ cluster specific background. First, we calculated the average C-to-U editing events in E-YTH^mut^ per cluster. Although ideally, we would like to remove background on a single cell level, this is not possible. Every single cell is unique, and a single cell in the E-YTH samples cannot be directly matched with any single cell from the E-YTH^mut^ to remove single cell specific background. Thus, instead, we calculated the average C-to-U editing events in E-YTH^mut^ per cluster. Therefore, for each individual cell in our E-YTH single cell data, we subsequently removed the average cluster specific background that each cell belongs to. We found that removing this m^6^A cluster specific background was essential to obtain accurate, cluster specific m^6^A characteristics.

### Comparison of bulk, single cell and m^6^A RNA immunoprecipitation datasets

The m^6^A overlap among single cell RNA-seq, bulk RNA-seq, and m^6^A-RIP was determined by identifying the intersection of genes where a m^6^A position (or region for m^6^A-RIP) was identified.

### Confirmation of differential m^6^A methylation

Four wild type mouse brain hippocampi were isolated and dissociated into single cells. OLG and IMC cells were isolated by FACS sorting using the Oligodendrocyte Marker O1 Monoclonal Antibody (O1), eFluor™ 660 (eBioscience, #50-6506-80) and the CX3CR1 Monoclonal Antibody (2A9-1), Alexa Fluor™ 488 (eBioscience, #53-6099-42), respectively. Following RNA isolation, m^6^A-RIP and reverse transcription, qPCR was performed with the following primers: *Colgalt1* (F:AAGAACTCAGATGTGCTCCAG; R:CTATAGTCCCAGGCAAGCAC), *Gsn* (F: CATCACAGTCGTTAGGCAGG; R:TGATGGCTTTGGTCCTTACTC). m^6^A-RIP versus matching input control samples were calculated.

### Identification of homogenous m^6^A sites in single cell sequencing data

To identify homogenous m^6^A sites, the m/k ratio was first calculated, as described above for bulk m^6^A sites. In addition, to identify truly homogenous m^6^A sites (m/k=1) in the single cell data, we excluded the possibility that any homogenous (m/k =1) sites might still represent mouse specific SNPs. Thus, we identified all m/k=1 m^6^A sites per cell for all cells, and then only kept those m/k=1 m^6^A sites that occurred in at least 5 other cells with a lower m/k ratio (m/k<0.9) and a minimum read overage of 10 per cell.

### Gene Ontology Analysis

Prior to gene ontology (GO) analysis, conserved homogenous m^6^A sites were identified. Such sites for each cell types consist of m/k=1 sites identified in our merged single cell RNA sequencing data that occur with an m/k<0.9 ratio in at least 5 cells of other cell types. For each cell type, GO biological process, molecular function and cellular component enrichment analyses were carried out on the set of genes where these conserved homogenous sites occur, using cluster Profiler (version 3.14.3) (Yu et al. 2012). Terms with an adjusted q-value<0.05 and p-value<0.05 were considered statistically significant. The top 5 GO terms were plotted, in addition to 5 selected terms that are statistically significant. Additionally, disease enrichment analysis was conducted on the human orthologue genes corresponding to the genes with conserved homogenous sites, using DOSE (version 3.12.0) and DisGeNET (Yu et al. 2015; Piñero et al. 2017). Terms with an adjusted q-value<0.05 and p-value<0.05 were considered statistically significant and plotted. Prism (V9.5.1) was used to plot any GO terms. The Rich factor is the ratio of gene numbers with m/k=1 m^6^A in a pathway term to all gene numbers annotated in this pathway term.

## DATA ACCESS

All raw and processed sequencing data generated in this study have been submitted to the NCBI Gene Expression Omnibus (GEO; https://www.ncbi.nlm.nih.gov/geo/) under accession number GSE240863.

Codes used in this study are available through GitHub (https://github.com/KoziolLaboratory/sc-m6a-hippocampus) and can be found in the Supplemental Code. Single cell RNA and m^6^A density UMAP visualizations can also be accessed *via* our interactive website (website: http://dart.cibr.ac.cn) and are provided as the Supplemental Material S1 & S2, respectively.

## ACKNOWLEDGMENTS

We are grateful to all members of the Koziol laboratory for valuable discussions and critical comments. We would like to thank the Genomics Center, Vector Core, Optical Imaging, HPC Facility, the Laboratory Animal Resource and Mass Spectrometry Center Facilities at the Chinese Institute for Brain Research for their generous support. We would like to specifically thank Tong Guo and Wenlong Li from our Laboratory Animal Resource Facility who supported us during the review process of this manuscript. We are thankful to Minmin Luo’s laboratory for their pAAV-CAG-EGFP-WPRE-SV40 plasmid. We are grateful to Yuyu Sheng from the BD Biosciences company for technical support. This project was supported by the Chinese Institute for Brain Research core grant, the Chinese Academy of Medical Sciences Innovation Fund for Medical Sciences (2019-I2M-5-015) and by the Beijing Natural Science Foundation (IS23091). M.T.A. contribution towards this work was funded by the Koziol lab. Y.P. and E.L. are supported by the Beijing Postdoctoral Research Foundation Fellowship (2020-YJ-002 and 2020- YJ-004, respectively) and by the International Postdoctoral Exchange Fellowship Program. We are grateful to the Chinese Institute for Brain Research Institute community for their generous support.

## Author contributions

F.S. designed and performed experiments, developed ideas, analyzed the data, supervised all experiments, assembled all data and wrote the paper. M.T.A. performed the bioinformatic analyses, critically evaluated data and reviewed the paper. Y.Z. helped with all animal experiments. Y.P. performed all UHPLC-MS/MS experiments and analyses. E.L. contributed through discussions, ideas and reviewed the paper. H.Z. developed the single cell m^6^A website. H.Z. was supported by L.Z. L.X. supported experimental and mouse work. M.J.K. conceived the study and designed experiments, analyzed the data, wrote the paper and supervised all research.

Supplemental Figures S1 to S8. (separate files)

Supplemental Tables S1 to S14. (separate files)

Supplemental Materials S1 to S3. (separate files)

Supplemental Code. (separate file)

