## Supplemental_Fig_S1 for "Single cell discovery of m^6^A RNA modifications in the hippocampus"

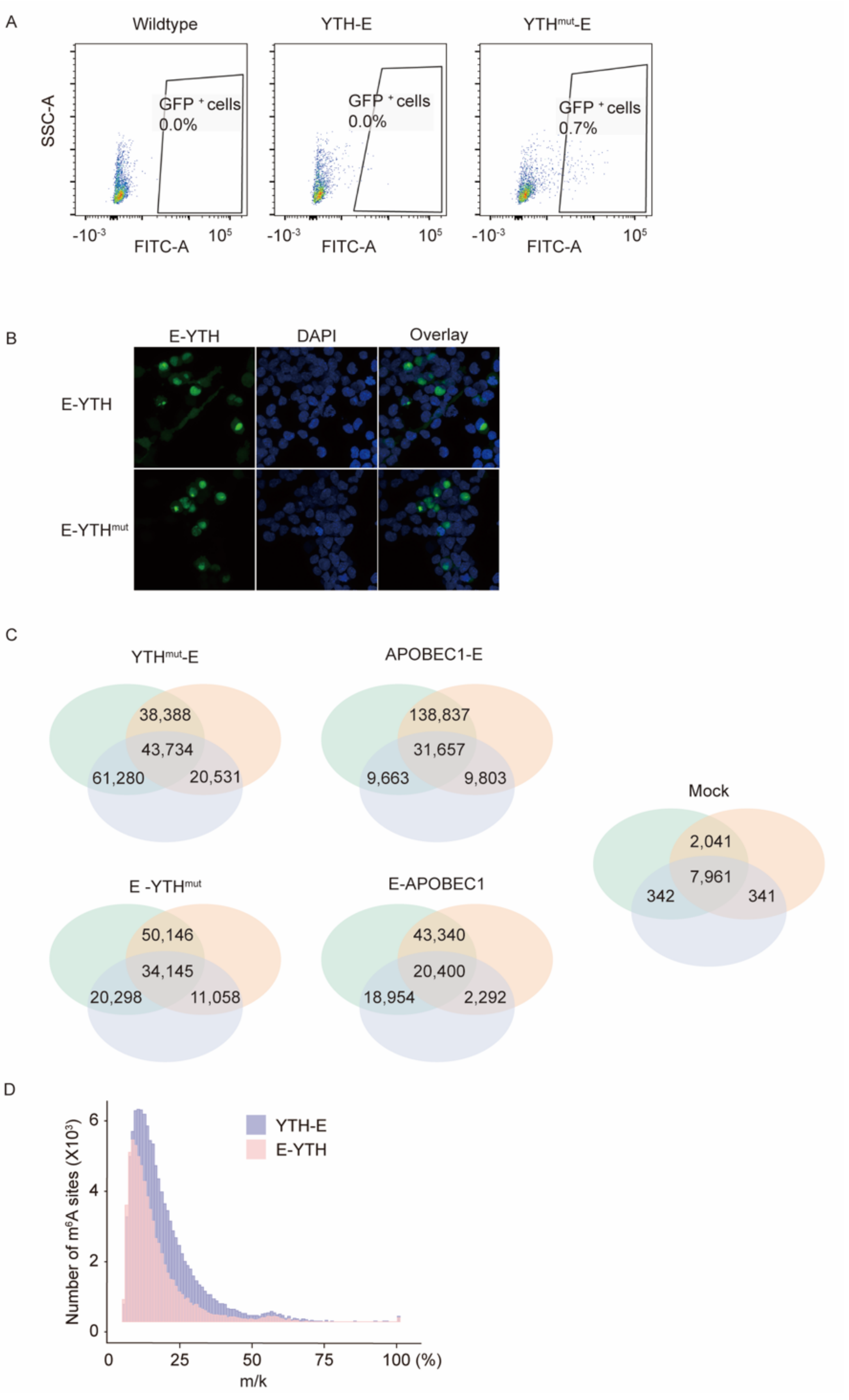

A
Wildtype
YTH-E
YTHmut-E
SSC-A
GFP + cells 0.0%
GFP + cells 0.0%
GFP + cells 0.7%
-10-3
FITC-A
105
B
E-YTH
DAPI
Overlay
E-YTH
E-YTHmut
C
YTHmut-E
38,388
43,734
61,280
20,531
APOBEC1-E
138,837
31,657
9,663
9,803
Mock
2,041
7,961
342
341
E -YTHmut
50,146
34,145
20,298
11,058
E-APOBEC1
43,340
20,400
18,954
2,292
D
YTH-E
E-YTH
Number of m6A sites (X103)
0
2
4
6
0
25
50
75
100 (%)
m/k

**Supplemental Fig S1. Improved $m^6A$ DART-seq detection in HEK293T cells.**

(A) Detection of EGFP positive cells in YTH-E, $YTH^{mut}$-E and wild type cells from mouse hippocampi. FACS gating is shown. The value indicates percentage of EGFP positive cells divided by total cells, excluding debris (P1 gate).

(B) Left: Confocal images of HEK293T cells transfected with E-YTH and E-$YTH^{mut}$. Representative images are shown, 24 hrs after transfection. Scale bar, 20 $\mu$m.

(C) Number of C-to-U editing events identified in each bulk HEK293T cell replicate for $YTH^{mut}$-E, E-$YTH^{mut}$, APOBEC1-E, E-POBEC1 and mock. Except for mock, the data was obtained following transfection and EGFP FACS sorting. n=3, Rep: separately cultured replicate.

(D) Histogram for YTH-E and E-YTH showing $m^6A$ counts over mutation per read (m/k) ratio. For both samples, background C-to-U editing sites identified in the in the corresponding control groups were removed. A minimum threshold of 5% was applied.
