## Supplemental_Fig_S2 for "Single cell discovery of m^6^A RNA modifications in the hippocampus"

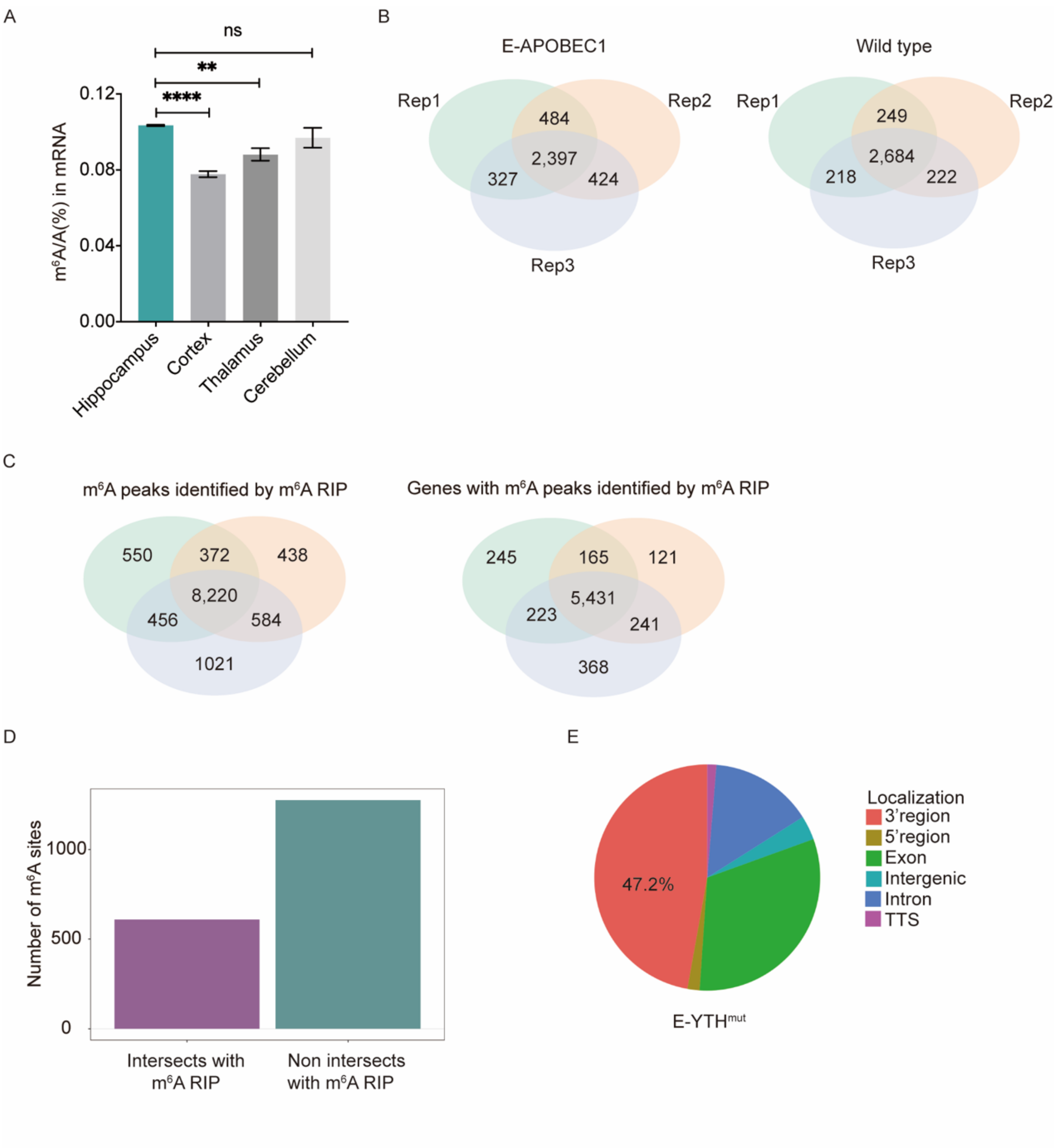


**Supplemental Fig S2. Detection of m$^6$A with bulk RNA-seq in mouse hippocampus.**

(A) LC-MS/MS quantification of m$^6$A within mRNA relative to unmodified adenosines in mRNA. mRNA was isolated from the hippocampus, cortex, thalamus and cerebellum of 3-month-old mice. n=3.

(B) Number of C-to-U editing events identified in each bulk RNA-seq replicate for controls including E-APOBEC1 and wild type mice. The data was obtained following AAV infection and EGFP FACS sorting. n=3, Rep: biological replicate, hippocampus from one mouse.

(C) Number of m$^6$A peaks (left) and number of genes with m$^6$A peaks (right) identified in each m$^6$A RIP replicate from wild type mouse hippocampi. n=3, Rep: biological replicate, hippocampus from one mouse.

(D) Histogram showing number of m$^6$A sites detected by bulk E-YTH that do and do not intersect with m$^6$A peaks identified with m$^6$A RIP.

(E) Pie chart showing C-to-U edit localization identified with E-YTH$^{mut}$ in mouse hippocampus. TTS: transcription termination site.
