## Supplemental_Fig_S3 for "Single cell discovery of m^6^A RNA modifications in the hippocampus"

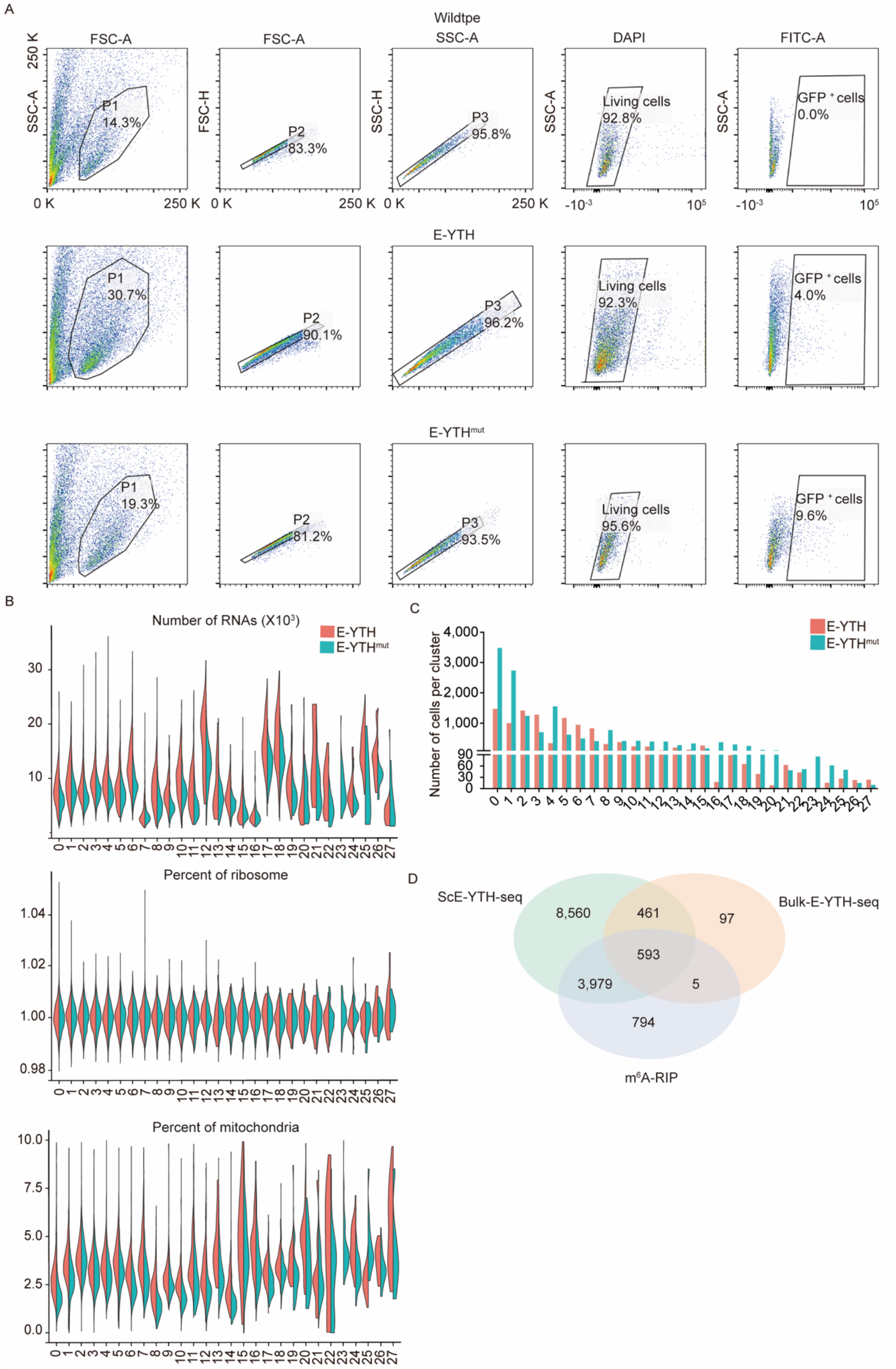
A
Wildtpe
E-YTH
E-YTHmut
FSC-A
SSC-A
DAPI
FITC-A
FSC-H
SSC-H
P1 14.3%
P2 83.3%
P3 95.8%
Living cells 92.8%
GFP + cells 0.0%
P1 30.7%
P2 90.1%
P3 96.2%
Living cells 92.3%
GFP + cells 4.0%
P1 19.3%
P2 81.2%
P3 93.5%
Living cells 95.6%
GFP + cells 9.6%
B
Number of RNAs (X10³)
Percent of ribosome
Percent of mitochondria
C
Number of cells per cluster
D
ScE-YTH-seq
Bulk-E-YTH-seq
m⁶A-RIP
8,560
461
97
593
3,979
5
794

**Supplemental Fig S3. Hippocampal single cell identification following AAV transduction.**

(A) Isolation of EGFP positive E-YTH, E-YTH$^{mut}$ and wild type cells from mouse hippocampi. FACS gating is shown. The value indicates percentage of cells in the selected gate.

(B) Single cell quality controls. Top: Violin plot showing the number of RNAs per cell cluster. Middle: Violin plot illustrating the percentage of ribosomal RNA per cell cluster. Bottom: Violin plot visualizing the percentage of mitochondrial RNA per cell cluster.

(C) Histogram with number of cells identified per cluster.

(D) Venn diagram showing the number of genes with m$^6$A regions detected by single cell, bulk and m$^6$A RIP RNA-seq.
