## Supplemental_Fig_S4 for "Single cell discovery of m^6^A RNA modifications in the hippocampus"

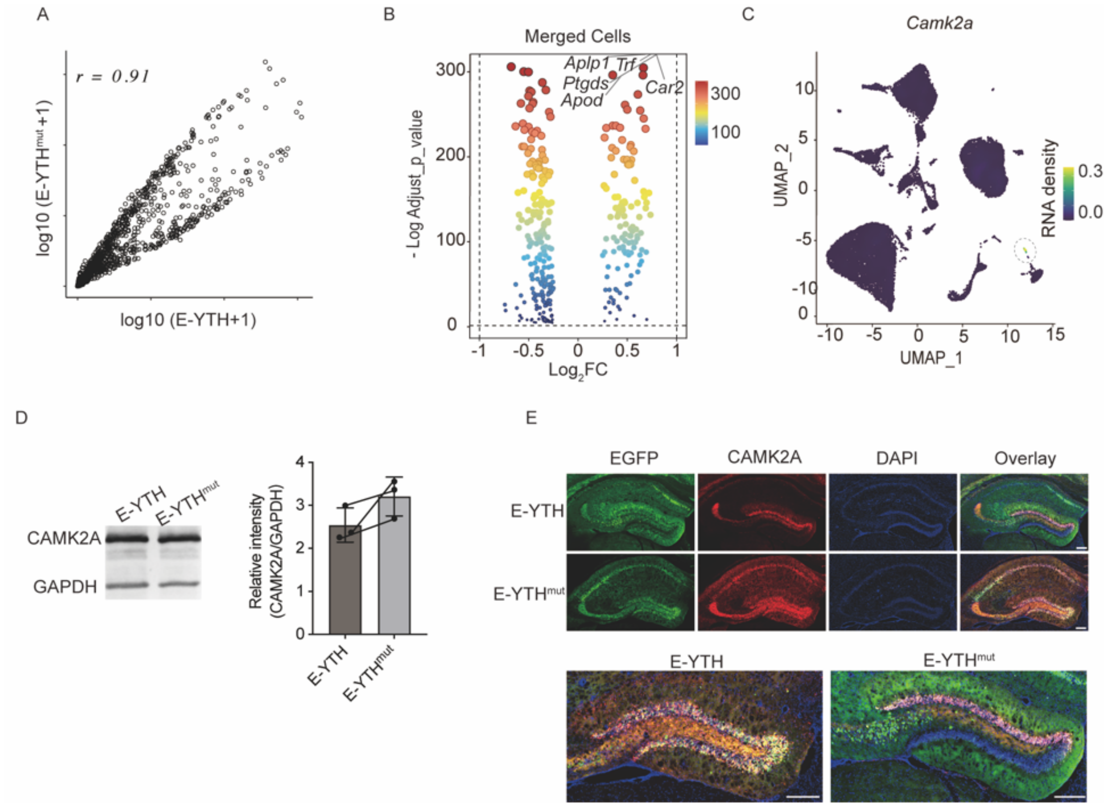


**Supplemental Fig S4. Transcriptional changes detected in E-YTH versus E-YTH$^{mut}$ single cell data.**

(A) Scatter plot between E-YTH and E-YTH$^{mut}$ single cell expression data (log10 TPM+1). Pearson correlation coefficient (*r*) is indicated. TPM: transcripts per million.

(B) Volcano plot comparing RNA levels of E-YTH and E-YTH$^{mut}$ single cell data. Single cell data was pooled and plotted to show all merged cells. Adjusted *p*-value<0.1 was applied before plotting. Adjusted *p*-value<0.05 and a Log2FC>1 and Log2FC<-1 are considered significant. Colour coded Legend value represents -Log10 *P* Value. FC: Fold change.

(C) Integrated UMAP (E-YTH and E-YTH$^{mut}$) with expression level for *Camk2a*. Legend colour represents RNA density. Circle was added to highlight region with increased RNA density.

(D) Western blot for CAMK2A and GAPDH. Hippocampi were processed following *E-Yth* or *E-Yth$^{mut}$* AAV injection. Normalized quantifications are shown. n=3 biological replicates.

(E) *Camk2a-mCherry* AAV was mixed with equal amounts of *Egfp* expressing E-YTH and E-YTH$^{mut}$ AAVs and the mix was subsequently injected into mouse hippocampi. Hippocampi were imaged to evaluate colocalization of CAMK2A and EGFP signals. CAMK2A expressing cells appear red, E-YTH and E-YTH$^{mut}$ expressing cells are green. Bottom image show increased zoom. In comparison to other AAV injection experiments, due to mixing with *Camk2a-mCherry* AAV, the amount of *E-Yth* and *E-Yth$^{mut}$* AAV viruses injected here is reduced by half. Scale bar: 200µm.
