## Supplemental_Fig_S5 for "Single cell discovery of m^6^A RNA modifications in the hippocampus"

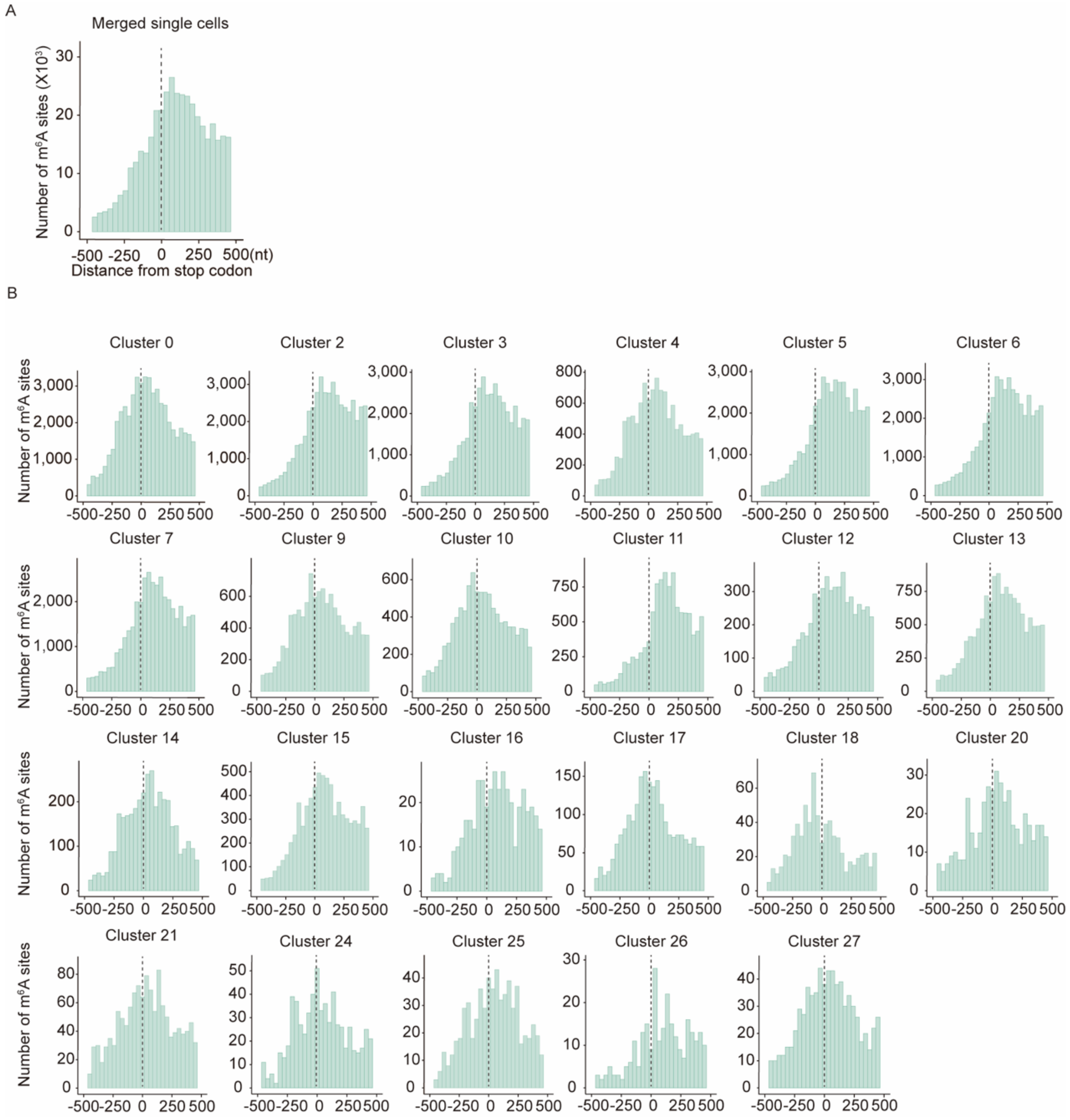


**Supplemental Fig S5. Hippocampal single cell metagene analysis surrounding the stop codon.**

(A) Metagene analysis of all cells merged identified by single cell sequencing. $m^6A$ density surrounding the stop codon (position 0). $m^6A$ sites were obtained after eliminating background editing sites.

(B) Metagene analysis of individual cell clusters identified by single cell sequencing. Clusters 0 through 7, 9 through 21 and 24 through 27 are shown. $m^6A$ density surrounding the stop codon (position 0). $m^6A$ sites were obtained after eliminating background editing sites.
