## Supplemental_Fig_S6 for "Single cell discovery of m^6^A RNA modifications in the hippocampus"

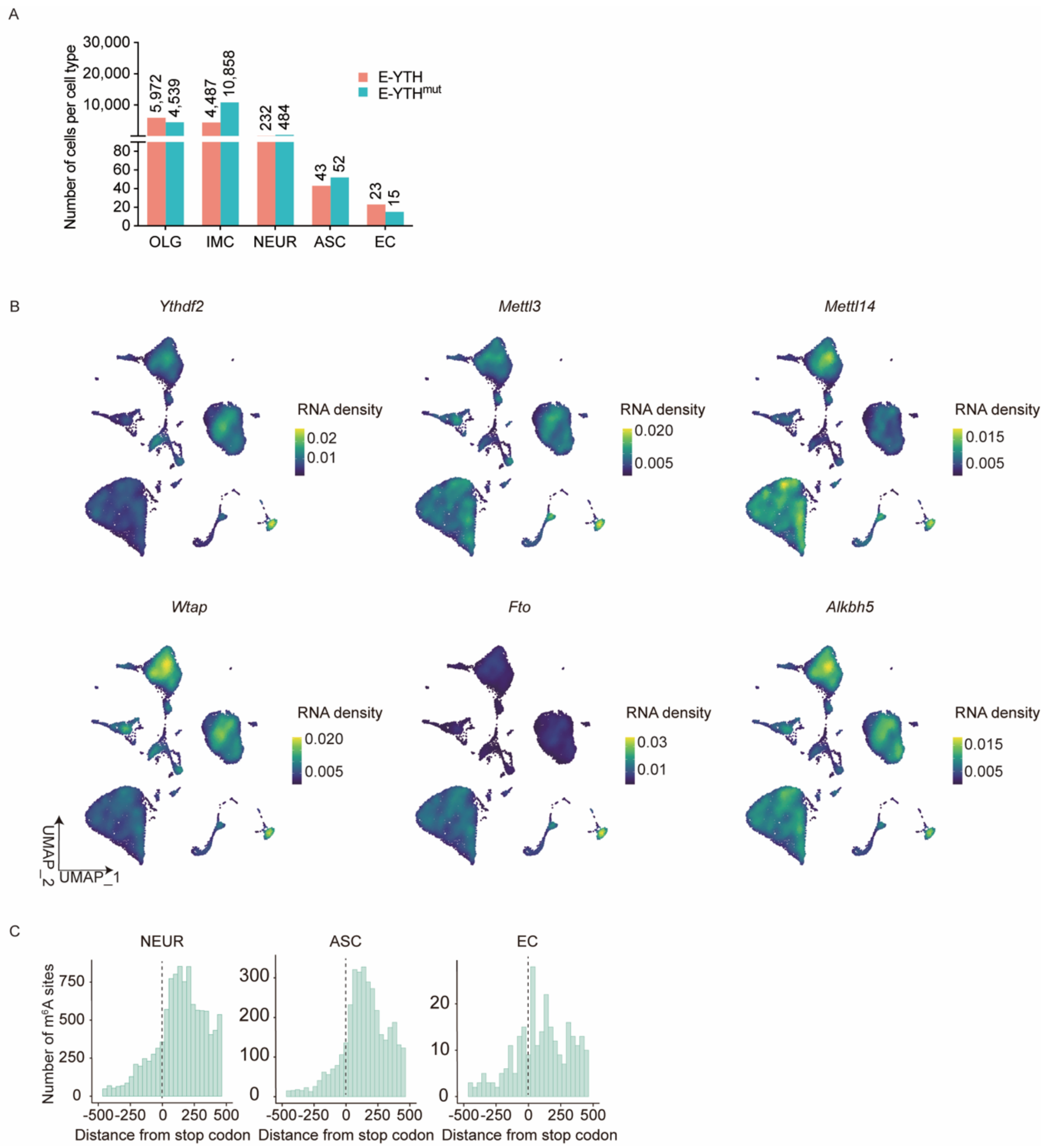


**Supplemental Fig S6. Hippocampal m$^6$A characteristics.**

(A) Histogram showing cell type numbers identified. OLG: oligodendrocyte cell lineage; NEUR: neuronal cell lineage; EC: endothelial cell lineage; IMC: immune cell lineage; ASC: astrocyte cell lineage.

(B) UMAP plot of single cell gene expression of *Ythdf2*, *Mettl3*, *Mettl14* and *Wtap* encoding m$^6$A methylases and of *Fto* and *Alkbh5* encoding m$^6$A demethylases. Legend colour represents RNA density.

(C) Distribution of m$^6$A surrounding the stop codon (0nt) identified by single cell sequencing for 3 main cell types. NEUR: neuronal cell lineage; EC: endothelial cell lineage; ASC: astrocyte cell lineage.
