## Supplemental_Fig_S7 for "Single cell discovery of m^6^A RNA modifications in the hippocampus"

A

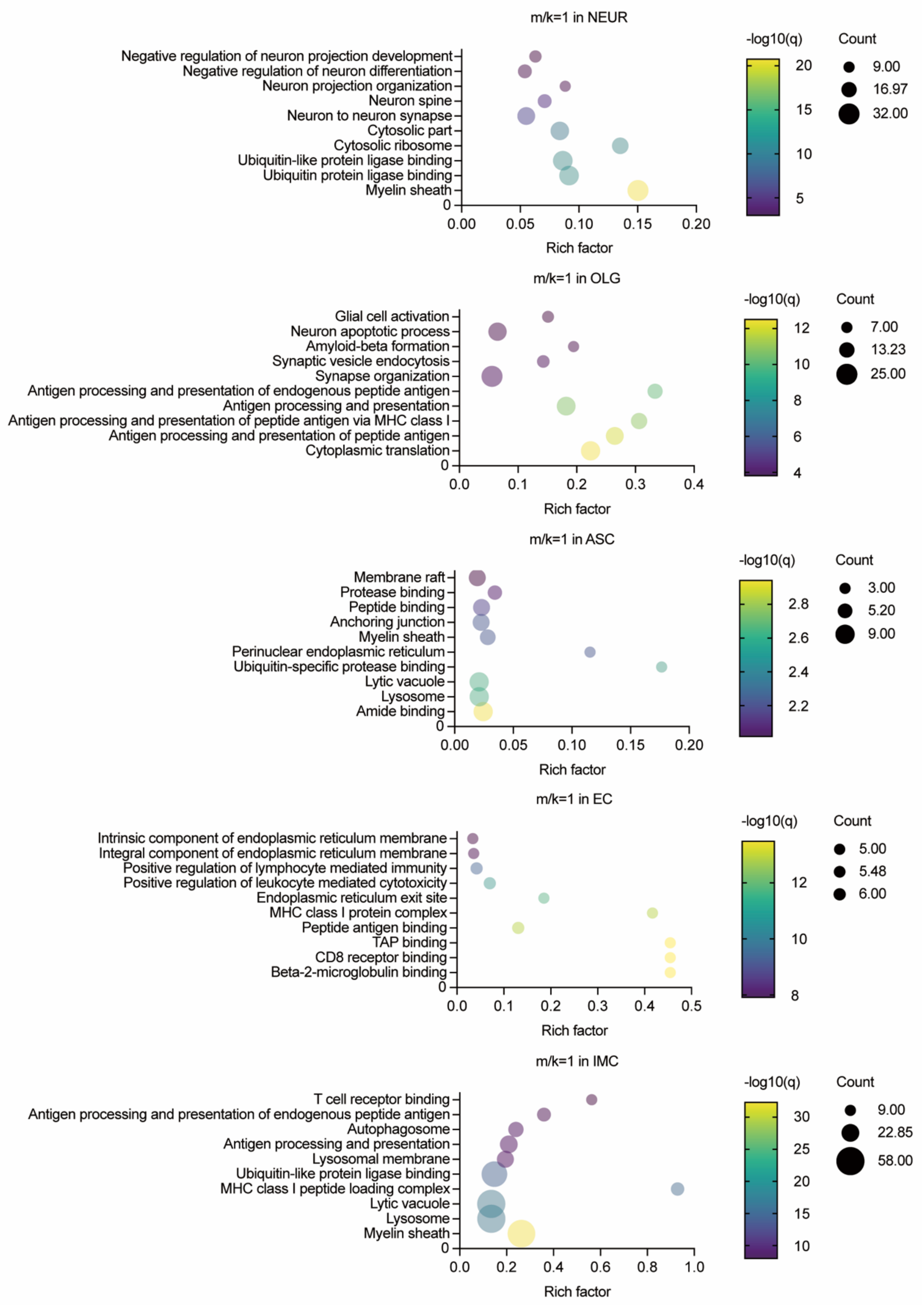
m/k=1 in NEUR
Negative regulation of neuron projection development
Negative regulation of neuron differentiation
Neuron projection organization
Neuron spine
Neuron to neuron synapse
Cytosolic part
Cytosolic ribosome
Ubiquitin-like protein ligase binding
Ubiquitin protein ligase binding
Myelin sheath
Rich factor
-log10(q)
Count
9.00
16.97
32.00
m/k=1 in OLG
Glial cell activation
Neuron apoptotic process
Amyloid-beta formation
Synaptic vesicle endocytosis
Synapse organization
Antigen processing and presentation of endogenous peptide antigen
Antigen processing and presentation
Antigen processing and presentation of peptide antigen via MHC class I
Antigen processing and presentation of peptide antigen
Cytoplasmic translation
Rich factor
-log10(q)
Count
7.00
13.23
25.00
m/k=1 in ASC
Membrane raft
Protease binding
Peptide binding
Anchoring junction
Myelin sheath
Perinuclear endoplasmic reticulum
Ubiquitin-specific protease binding
Lytic vacuole
Lysosome
Amide binding
Rich factor
-log10(q)
Count
3.00
5.20
9.00
m/k=1 in EC
Intrinsic component of endoplasmic reticulum membrane
Integral component of endoplasmic reticulum membrane
Positive regulation of lymphocyte mediated immunity
Positive regulation of leukocyte mediated cytotoxicity
Endoplasmic reticulum exit site
MHC class I protein complex
Peptide antigen binding
TAP binding
CD8 receptor binding
Beta-2-microglobulin binding
Rich factor
-log10(q)
Count
5.00
5.48
6.00
m/k=1 in IMC
T cell receptor binding
Antigen processing and presentation of endogenous peptide antigen
Autophagosome
Antigen processing and presentation
Lysosomal membrane
Ubiquitin-like protein ligase binding
MHC class I peptide loading complex
Lytic vacuole
Lysosome
Myelin sheath
Rich factor
-log10(q)
Count
9.00
22.85
58.00

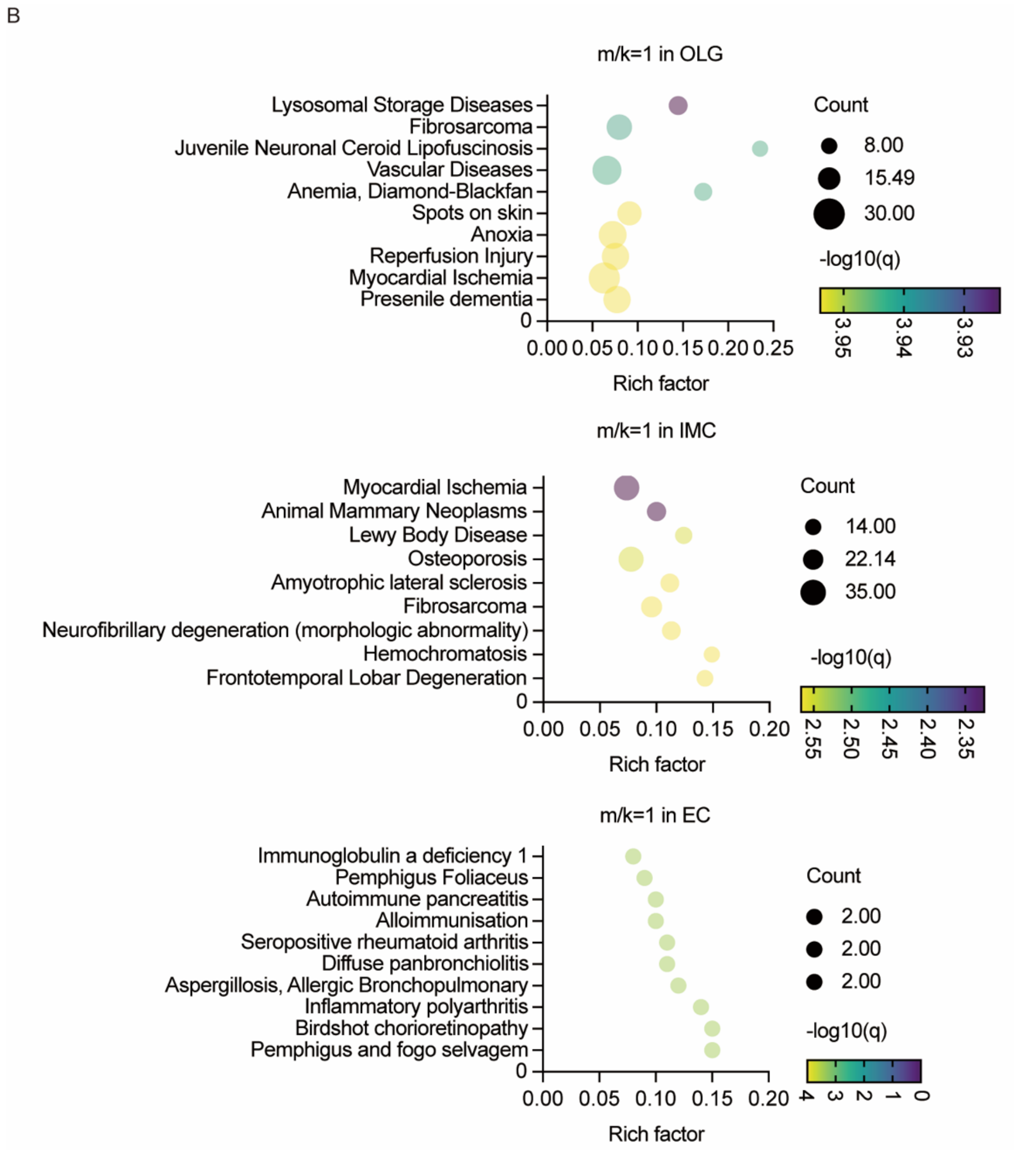


**Supplemental Fig S7. Gene Ontology Analyses of conserved homogenous m$^6$A sites.**

(A) Biological process, molecular function, and cellular component gene enrichment analyses (GO). Terms with an adjusted q-value<0.05 and p-value<0.05 were considered statistically significant. The top 5 GO terms were plotted for each cell lineage, in addition to 5 selected statistically significant terms. OLG: oligodendrocyte cell lineage; NEUR: neuronal cell lineage; EC: endothelial cell lineage; IMC: immune cell lineage; ASC: astrocyte cell lineage. The Rich factor represent the conserved homogenous m$^6$A site count divided by total counts.

(B) Disease gene enrichment analyses (GO). GO terms with an adjusted q-value<0.05 and p-value<0.05 are plotted for OLG, IMC, EC. OLG: oligodendrocyte cell lineage; EC: endothelial cell lineage; IMC: immune cell lineage.
