## Supplemental_Fig_S8 for "Single cell discovery of m^6^A RNA modifications in the hippocampus"

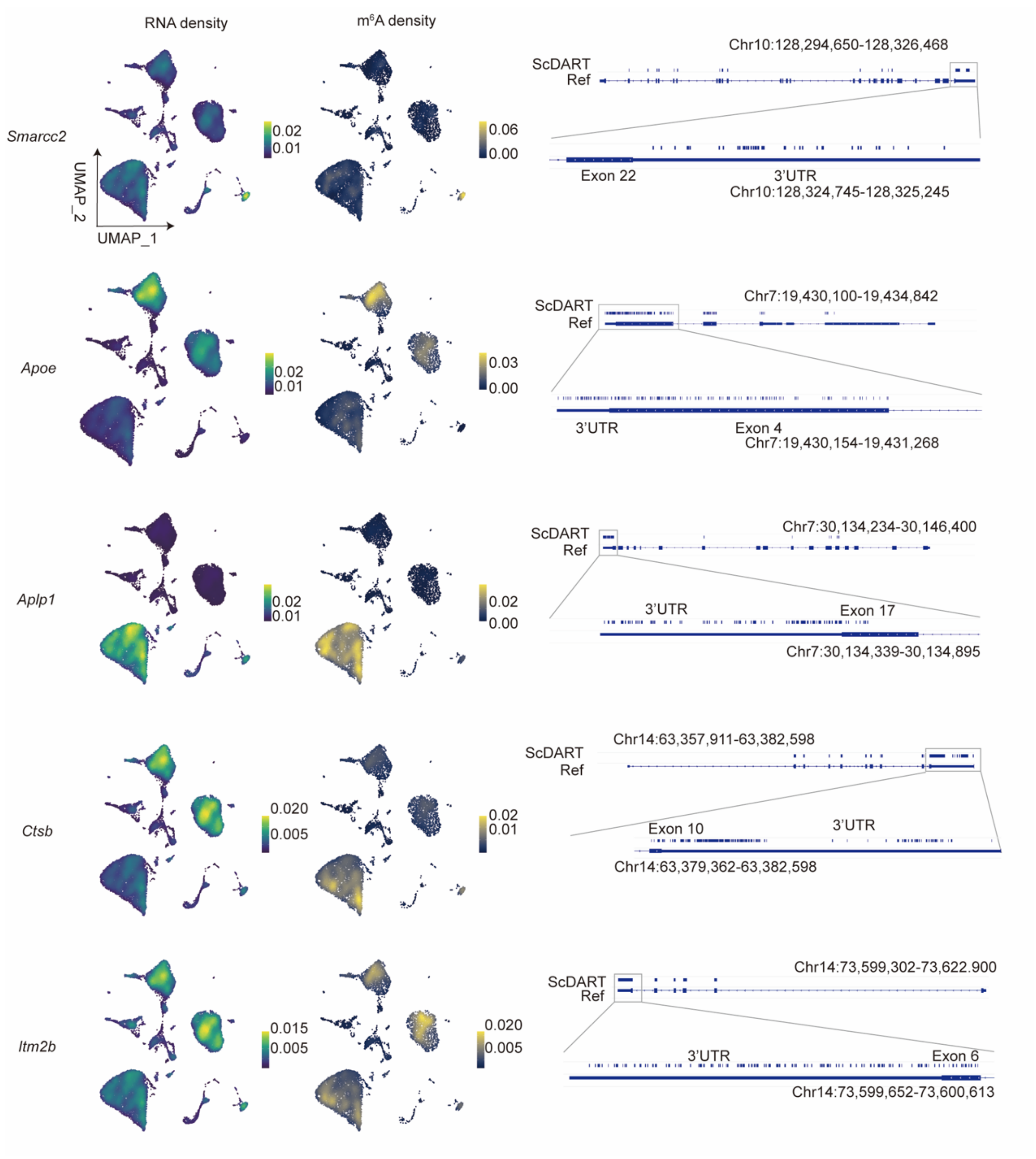

RNA density
m6A density
Smarcc2
UMAP_2
UMAP_1
0.02
0.01
0.06
0.00
ScDART
Ref
Chr10:128,294,650-128,326,468
Exon 22
3'UTR
Chr10:128,324,745-128,325,245
Apoe
0.02
0.01
0.03
0.00
Chr7:19,430,100-19,434,842
3'UTR
Exon 4
Chr7:19,430,154-19,431,268
Aplp1
0.02
0.01
0.02
0.00
Chr7:30,134,234-30,146,400
3'UTR
Exon 17
Chr7:30,134,339-30,134,895
Ctsb
0.020
0.005
0.02
0.01
Chr14:63,357,911-63,382,598
Exon 10
3'UTR
Chr14:63,379,362-63,382,598
Itm2b
0.015
0.005
0.020
0.005
Chr14:73,599,302-73,622.900
3'UTR
Exon 6
Chr14:73,599,652-73,600,613

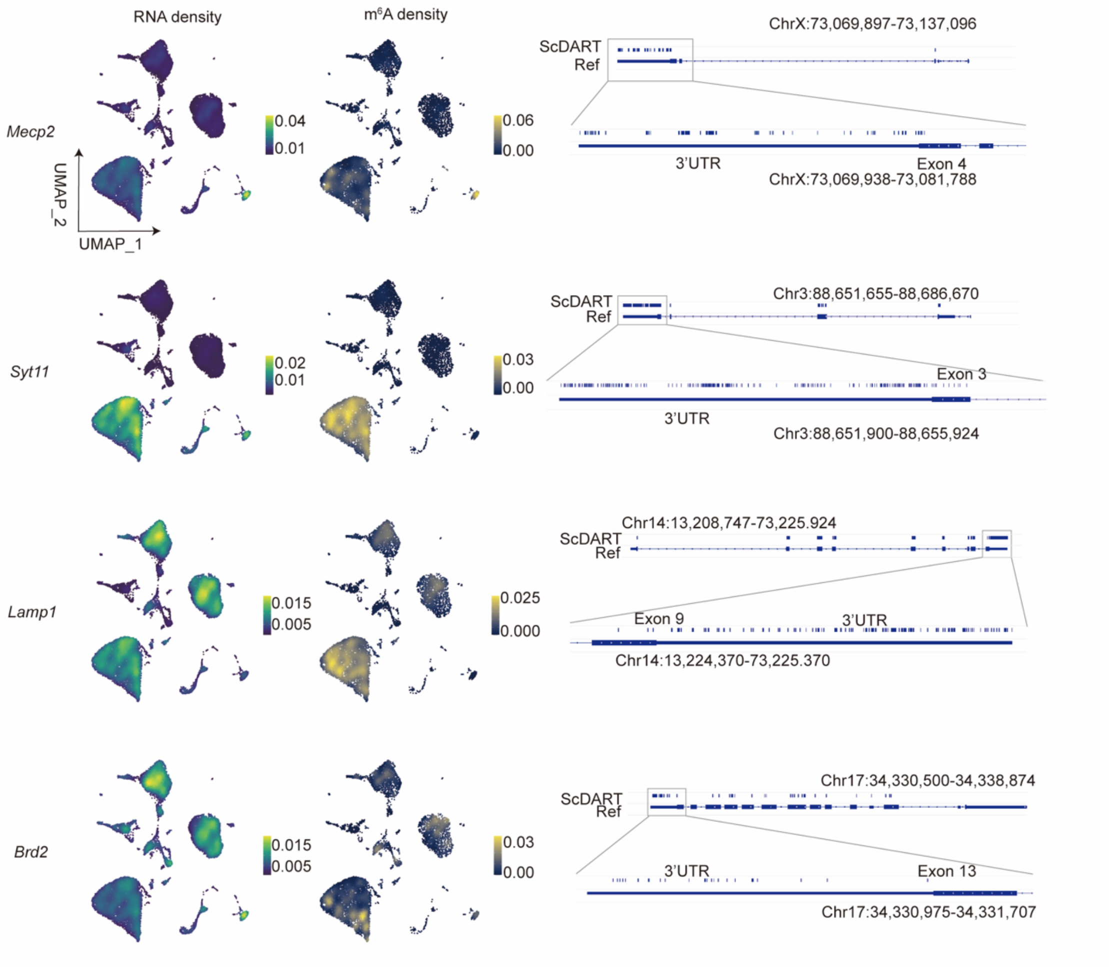


**Supplemental Fig S8. Hippocampal m$^6$A characteristics.**

Localizations of m$^6$A for different genes. Left: UMAP plot of RNA density for one gene per cell. Legend colour represents RNA density. Middle: UMAP plot of m$^6$A density for one gene per cell. Legend colour represents m$^6$A density on RNAs transcribed from one gene. Right: IGV RefSeq gene annotations with editing sites representing adjacent m$^6$A sites is shown. Last exon with 3'UTR region is illustrated with higher magnitude. Chr: Chromosome number; Units: base pairs.
